# Spatial alanine metabolism determines local growth dynamics of *Escherichia coli* colonies

**DOI:** 10.1101/2021.02.28.433255

**Authors:** Francisco Díaz-Pascual, Martin Lempp, Kazuki Nosho, Hannah Jeckel, Jeanyoung K. Jo, Konstantin Neuhaus, Raimo Hartmann, Eric Jelli, Mads Frederik Hansen, Alexa Price-Whelan, Lars E.P. Dietrich, Hannes Link, Knut Drescher

## Abstract

Bacteria commonly live in spatially structured biofilm assemblages, which are encased by an extracellular matrix. Metabolic activity of the cells inside biofilms causes gradients in local environmental conditions, which leads to the emergence of physiologically differentiated subpopulations. Information about the properties and spatial arrangement of such metabolic subpopulations, as well as their interaction strength and interaction length scales are lacking, even for model systems like *Escherichia coli* colony biofilms grown on agar-solidified media. Here, we use an unbiased approach, based on temporal and spatial transcriptome and metabolome data acquired during *E. coli* colony biofilm growth, to study the spatial organization of metabolism. We discovered that alanine displays a unique pattern among amino acids and that alanine metabolism is spatially and temporally heterogeneous. At the anoxic base of the colony, where carbon and nitrogen sources are abundant, cells secrete alanine *via* the transporter AlaE. In contrast, cells utilize alanine as a carbon and nitrogen source in the oxic nutrient-deprived region at the colony mid-height, *via* the enzymes DadA and DadX. This spatially structured alanine cross-feeding influences cellular viability and growth in the cross-feeding-dependent region, which shapes the overall colony morphology. More generally, our results on this precisely controllable biofilm model system demonstrate a remarkable spatiotemporal complexity of metabolism in biofilms. A better characterization of the spatiotemporal metabolic heterogeneities and dependencies is essential for understanding the physiology, architecture, and function of biofilms.

## Introduction

After bacterial cell division on surfaces, daughter cells often remain in close proximity to their mother cells. This process can yield closely packed populations with spatial structure, which are often held together by an extracellular matrix. Such spatially structured assemblages, called biofilms (Flemming et al., 2016), are estimated to be the most abundant form of microbial life on Earth (Flemming & Wuertz, 2019). The metabolic activity of cells inside these dense populations leads to spatial gradients of oxygen, carbon and nitrogen sources, as well as many other nutrients and waste products (Ackermann, 2015; Evans et al., 2020; Pacheco et al., 2019; Stewart & Franklin, 2008). Cells in different locations within these assemblages therefore inhabit distinct microenvironments. The physiological responses to these microenvironmental conditions result in spatially segregated subpopulations of cells with different metabolism (Barroso-Batista et al., 2020; D’Souza et al., 2018; Stewart & Franklin, 2008). Bacterial growth into densely-packed spatially structured communities, and metabolic activity of the constituent cells, therefore naturally lead to physiological differentiation (Evans et al., 2020; Røder et al., 2020; Serra, Richter, Klauck, et al., 2013; Stewart & Franklin, 2008).

Metabolic and phenotypic heterogeneities are frequently observed in multi-species communities (Garg et al., 2016; Henson et al., 2019; Kim et al., 2020; Pacheco et al., 2021) and in single-species biofilm populations (Cole et al., 2015; Dal Co et al., 2019; Lin et al., 2018; Moree et al., 2012; Rani et al., 2007; Serra, Richter, Klauck, et al., 2013; Teal et al., 2006). Identifying the origins of these heterogeneous subpopulations, and how they interact with each other, is important for understanding the development and function of biofilms (Cole et al., 2015; Lin et al., 2018; Liu et al., 2015; Prindle et al., 2015). Multi-species biofilms predominate in the environment, yet they are highly complex and feature many concurrent intra-species and inter-species interaction processes, which may be organized in space and time (Brislawn et al., 2019; Garg et al., 2016; Kim et al., 2020). Due to this complexity of multi-species biofilms, it is often difficult to disentangle whether spatiotemporal physiological differentiation or a particular interaction between subpopulations is caused by the relative position of the species, or the position of each species in the context of the entire community, or the mutual response of different species to each other. In contrast, single-species biofilms offer precisely controllable model systems with reduced complexity for understanding basic mechanisms of metabolic differentiation and the interaction of subpopulations in communities. Investigations of phenotypic heterogeneity in single-species assemblages have already revealed fundamental insights into metabolic interactions of subpopulations with consequences for the overall fitness and growth dynamics of the assemblage (Arjes et al., 2021; Cole et al., 2015; Evans et al., 2020; Lin et al., 2018; Liu et al., 2015, 2017; Wolfsberg et al., 2018). However, even for single-species biofilms, the extent of metabolic heterogeneity and metabolic dependencies of subpopulations are unclear.

To obtain an unbiased insight into the spatial organization of metabolism inside biofilms, we measured metabolome and transcriptome dynamics during the development of *E*. *coli* colony biofilms on a defined minimal medium that was solidified with agar. Our model system enabled highly reproducible colony growth and precise control of environmental conditions, which allowed us to detect phenotypic signatures of subpopulations. The temporally and spatially resolved data revealed that alanine metabolism displays a unique pattern during colony growth. We determined that secretion of alanine occurs in a part of the anaerobic region of the colony, where the carbon and nitrogen sources are abundant. The secreted alanine is then consumed in a part of the aerobic region of the colony, where glucose and ammonium from the minimal medium are lacking. This spatially organized alanine cross-feeding interaction occurs over a distance of tens of microns, and has important consequences for the viability and growth of the localized cross-feeding-dependent subpopulation, and for the global colony morphology.

## Results

### Colony growth transitions and global metabolic changes

To investigate the spatiotemporal organization of metabolism inside *E. coli* colonies, we first characterized the basic colony growth dynamics and morphology on solid M9 minimal medium, which contained glucose and ammonium as the sole carbon and nitrogen sources, respectively (Figure 1A). For all measurements performed on colonies, including microscopy-based measurements, the colonies were grown on top of filter membranes that were placed on M9 agar (Figure 1—figure supplement 1A). These membranes enabled rapid sampling of the colonies for transcriptome and metabolome analyses. When we grew *E. coli* wild type colonies in these conditions, we observed motility on the filter membranes on M9 agar, which led to heterogeneous colony development and an asymmetric colony morphology (Figure 1—figure supplement 1B). However, to study the spatiotemporal metabolism in *E. coli* colonies as precisely as possible, and to distinguish even small phenotypic differences between strains and conditions, we required a colony growth behavior that is as reproducible as possible. Therefore, a strain lacking flagella due to the deletion of the flagellin (Δ*fliC*) was used as the parental strain for all following experiments, which resulted in highly reproducible and axially symmetric colony morphologies (Figure 1—figure supplement 1C). Colonies generally displayed two growth phases: colonies showed exponential growth in volume, height, and diameter for up to ~24 h, followed by linear growth (Figure 1A,B). This transition in colony growth dynamics has previously been observed, and was hypothesized to be caused by a change in metabolism due to altered nutrient penetration and consumption for colonies above a certain size (Pirt, 1967; Warren et al., 2019).

**Figure 1.**
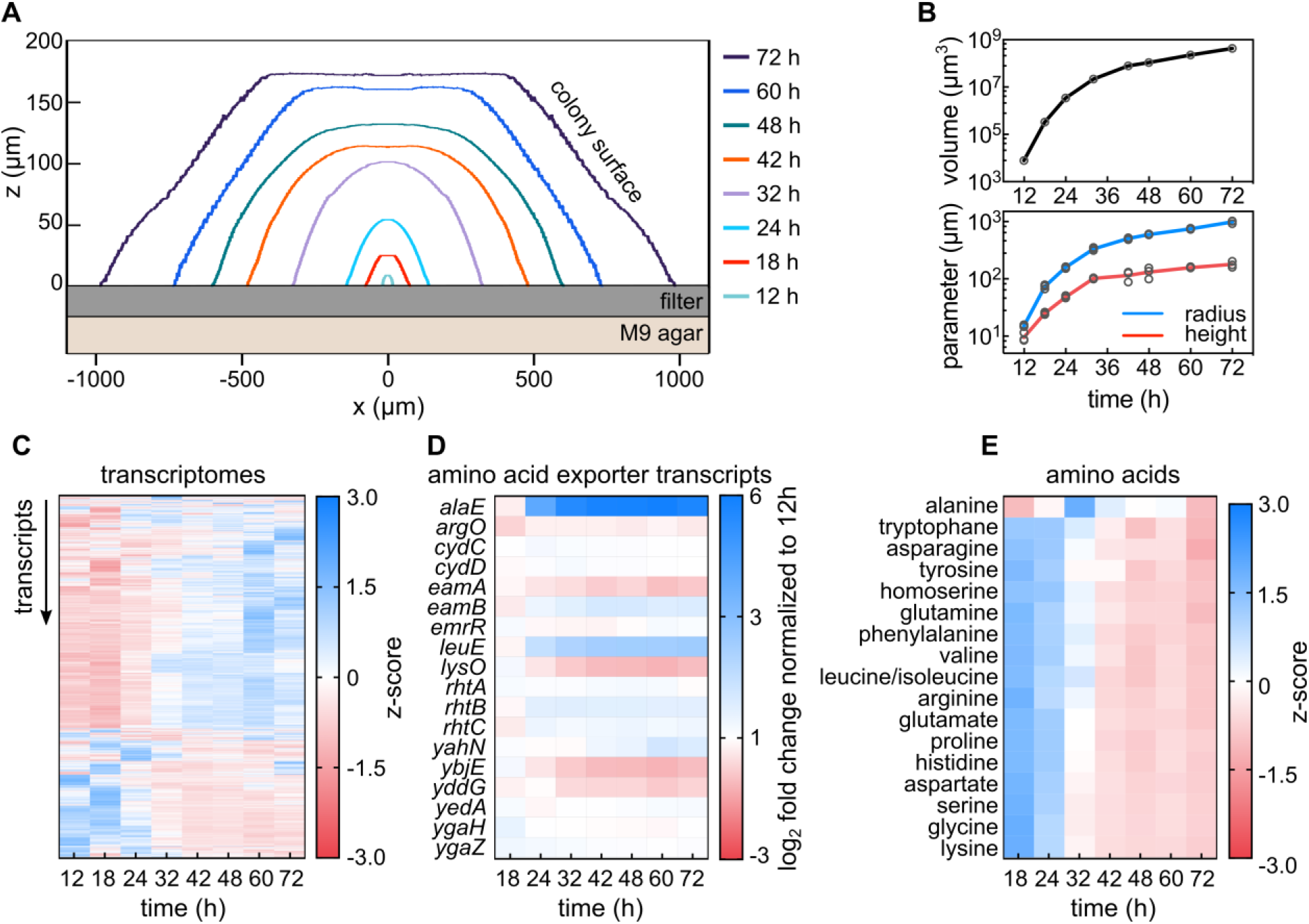
Transcriptomes and metabolomes during *E. coli* colony biofilm growth reveal metabolic transition and a unique role of alanine. (**A**) Cross-sectional profiles of representative *E. coli* colonies grown on membrane filters placed on solid M9 agar at different time points. (**B**) Volume (top panel), radius and height (bottom panel) of colonies as a function of time indicate a growth rate transition around 24-32 h. All replicates are shown as individual data points, with a line connecting the mean values, *n* = 3. (**C**) Dynamics of expression profiles of 4,231 genes during colony growth reveal physiological transition between 24-32 h, *n* = 3. (**D**) Mean expression fold-changes of known amino acid exporters during colony growth, calculated in comparison to 12 h colonies, *n* = 3. (**E**) Comparison of amino acid profiles in whole colonies, measured with mass spectrometry during colony growth; *n* = 3. Levels of amino acids are shown in Figure 1—figure supplement 5.

To further characterize the phenotypic changes that occur during colony growth, we performed time-resolved whole-colony transcriptome measurements (Figure 1C and Figure 1—figure supplement 2A). The transcriptomes revealed a major change after 24 h of growth (Figure 1C), which reflects the change in growth dynamics that was apparent in the morphological parameters (Figure 1B). After 72 h of growth, 966 out of 4,231 detected genes were differentially expressed in comparison to the 12 h time point (log_2_-fold-changes >1 or <−1, and FDR-adjusted *p*-value <0.05). The whole-colony transcriptomes also showed that biofilm matrix biosynthesis genes are expressed continuously during colony growth (Figure 1—figure supplement 3). Furthermore, the transcriptomes confirmed that well-known pathways such as mixed acid fermentation and tricarboxylic acid (TCA) cycle were differentially regulated after 24 h of growth (Figure 1—figure supplement 4A,C). These results are consistent with the hypothesis that above a certain colony size, which is reached between 24-32 h in our conditions, the consumption of oxygen by cells in the outer region of the colony causes a large anoxic region inside the colony.

Although it is commonly assumed that *E. coli* subpopulations cross-feed acetate (Cole et al., 2015; Dal Co et al., 2019; San Roman & Wagner, 2020; Wolfsberg et al., 2018), the temporal transcriptomes did not reveal strong regulation of acetate metabolism transcripts during the major transition in colony metabolism described above (Figure 1—figure supplement 4A). Lactate, formate, and succinate biosynthesis, however, displayed interesting dynamics during colony growth (Figure 1— figure supplement 4A). Apart from these metabolites, we noticed peculiar patterns in the expression of amino acid pathways. The gene expression levels for some amino acid transporters remained unchanged during colony growth, while a few were >2-fold up- or down-regulated after 24 h or 32 h of growth (Figure 1D). Interestingly, the gene coding for the alanine exporter *alaE* (Hori, Yoneyama, et al., 2011) showed strong expression changes (~50-fold increase) in 72-h-colonies relative to 12-h-colonies (Figure 1D).

To further characterize the observed metabolic transition between 24-32 h of colony growth, we measured the amino acid profiles of the colonies using mass spectrometry and normalized them by the colony biomass. To obtain sufficient biomass for these measurements, the colonies needed to be grown for at least 18 h. All amino acid abundances decreased during colony growth – except for alanine, which remained relatively constant with a peak abundance at 32 h (Figure 1E, Figure 1—figure supplement 2B, and Figure 1—figure supplement 5). Thus, both transcriptome and metabolome data suggest a unique change of alanine metabolism during colony growth, which led us to investigate its role in *E. coli* colonies.

### Spatial regulation of alanine transport and degradation

To test whether alanine metabolism is spatially heterogeneous during biofilm growth, we developed a method to measure transcriptomes with spatial resolution in the colonies. The method is based on the oxygen-dependence of chromophore maturation of the fluorescent protein mRuby2 (Lam et al., 2012; Tsien, 1998). Using a strain that constitutively expresses mRuby2 from a chromosomal locus, so that all cells in the colony produce mRuby2, we observed that only the air-facing region of the colony was fluorescent in colonies that had grown for 72 hours. Quantification of the mRuby2 fluorescence showed a slightly steeper decrease of fluorescence in the horizontal *xy*-direction into the colony, compared with the vertical *z*-direction (Figure 2A). By using an oxygen microsensor to directly measure oxygen levels inside the colony in the *z*-direction (Jo et al., 2017), we determined that the mRuby2 fluorescence was a reliable indicator of oxygen levels (Figure 2A). During colony growth, the fraction of fluorescent cells in the colony decreased (insert in Figure 2B), ultimately leading to a thin layer of fluorescent cells in the air-facing part of the colony (Figure 2B).

**Figure 2.**
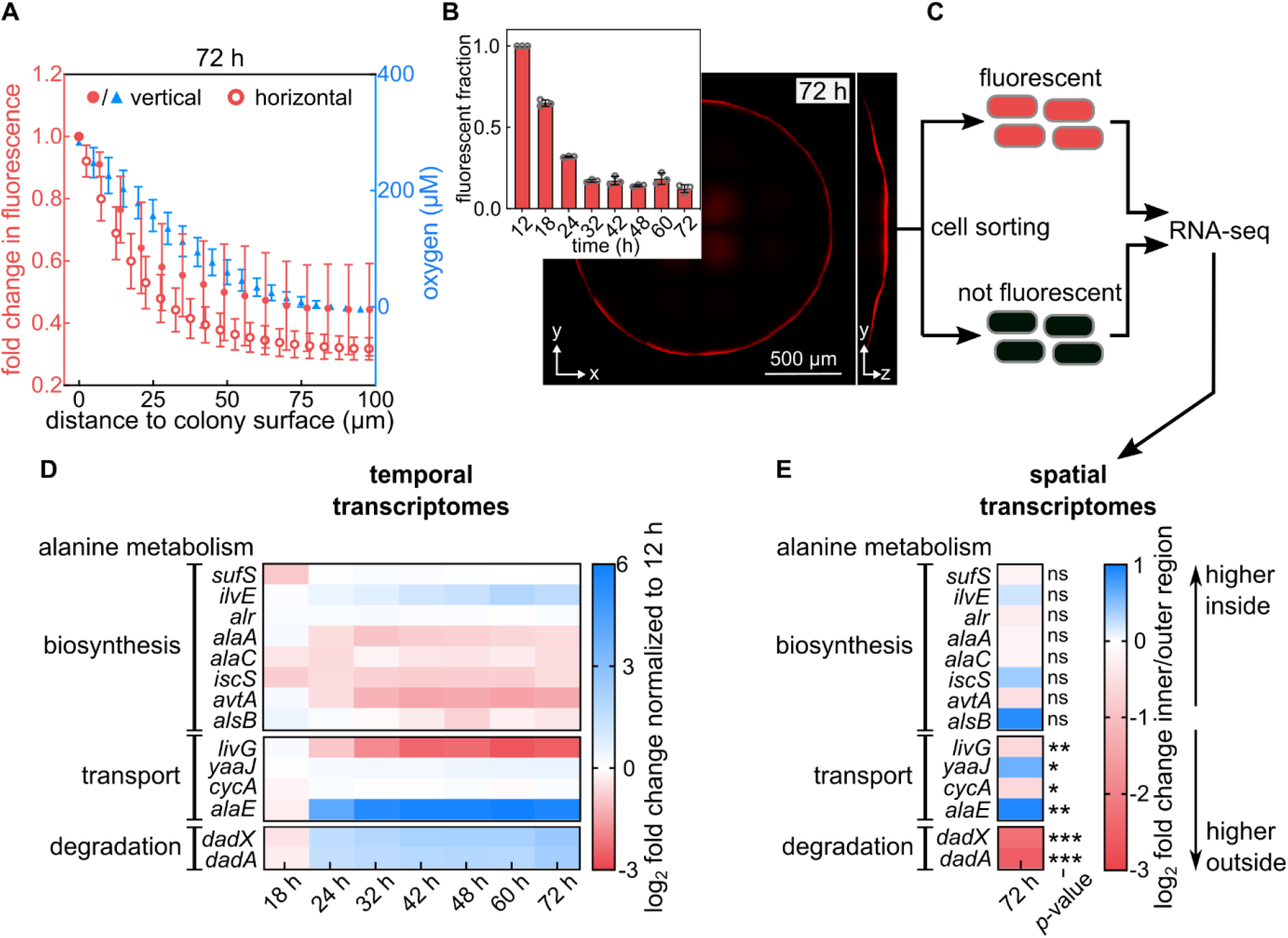
Alanine transport and degradation are spatially regulated within colony biofilms. (**A**) Measurement of oxygen penetration into colonies grown for 72 h on filter membranes on M9 agar using two different methods (for strain KDE722). Left axis (red): Intensity of mRuby2 fluorescence within a vertical cylinder of radius 3.3 μm in the center of the colony (filled circles), and intensity of mRuby2 fluorescence within a horizontal plane at the base of the colony (open circles, mean ± s.d., *n* = 3). Right axis (blue): Direct measurement of oxygen levels acquired by vertical scanning of an oxygen microsensor at the center of the colony (blue triangles, mean ± s.d., *n* = 10). (**B**) Confocal image of a representative 72 h colony of a strain that constitutively expresses mRuby2 (KDE722). Insert: Fraction of fluorescent biovolume inside colonies grown for different times. Data are mean ± s.d., *n* = 3. (**C**) Scheme of the sorting procedure: Cells from a 72 h colony are separated, fixed using formaldehyde, and sorted according to their mRuby2 fluorescence, followed by RNA-seq, as described in the methods section. (**D**) Heatmaps showing the mean fold-change (*n* = 3) of expression levels of genes involved in alanine metabolism, from whole-colony measurements. Fold-changes are computed relative to the 12 h timepoint. (**E**) Spatial transcriptome results, quantified as fold-changes between the inner region (no mRuby2 fluorescence) and outer region (high mRuby2 fluorescence) regions of colonies grown for 72 h. Blue color in the heatmap indicates genes with higher transcript levels in the inner region of the colony, red color indicates higher transcripts in the outer region of the colony. Data are means, *n* = 4. Non-significant differences between spatial regions are labelled “ns”. *p*-values correspond to false discovery rate (FDR)-adjusted *p*-values: * *p* < 0.05, ** *p* < 0.01, *** *p* < 0.001.

To eliminate the possibility that the observed mRuby2 fluorescence profile was due to imaging artefacts, such as insufficient laser penetration into the colony, we disrupted 72-h-colonies and imaged the resulting, well-separated single cells. The time between colony disruption and imaging for this control experiment was less than 2 min, which is substantially shorter than the mRuby2 fluorophore maturation time (~150 min (Lam et al., 2012)). In these images, only some cells displayed fluorescence (Figure 2—figure supplement 1A), indicating that the fluorescence gradient we observed in Figure 2B is not an imaging artefact. We further reasoned that, because the oxygen gradient in the colony is created by oxygen consumption by cells in the air-facing region (Klementiev et al., 2020; Stewart & Franklin, 2008), the oxygen gradient (and therefore the mRuby2 fluorescence gradient) should disappear when metabolic processes that consume oxygen are prevented. To test this, we starved the colonies by moving the filter membrane carrying the colonies to an M9 agar plate lacking glucose. We observed that in this case, mRuby2 proteins that are located in the formerly dark anaerobic region became fluorescent. This finding is consistent with the interpretation that in the absence of glucose, cells consume less oxygen so that molecular oxygen can penetrate into the colony to enable chromophore maturation of mRuby2 in for formerly anoxic region (Figure 2—figure supplement 1B,C). Together, these control experiments and the direct measurements of oxygen levels show that the difference in fluorescence levels in our system reflect the spatial position of the cells in the colony.

To obtain spatial transcriptomes, we then separated colonies grown for 72 h into individual cells, immediately fixed them with formaldehyde (which prevents mRuby2 fluorophore maturation in the presence of oxygen without altering the transcriptomes; Figure 2—figure supplement 2), and subjected them to fluorescence-activated cell sorting (FACS) to separate the aerobic (fluorescent) and anaerobic (not fluorescent) populations (Figure 2C and Figure 2—figure supplement 3). The resulting two cell populations were then analyzed using RNA-seq. The spatial transcriptome comparison showed that, as expected, genes involved in the TCA cycle and mixed acid fermentation were differentially expressed between the inner and outer regions of the colony (Figure 1—figure supplement 4B,D). This result demonstrates that the method successfully separated the fermenting population inside the colonies from the respiring population in the outer layer, which serves as a qualitative verification of the experimental methodology.

In both the spatially and temporally resolved transcriptomes, we observed changes in alanine transport and degradation, but not in alanine biosynthesis (Figure 2D,E; a schematic diagram of alanine metabolism pathways is shown in Figure 2—figure supplement 4). In particular, the expression of the alanine exporter gene *alaE* was significantly up-regulated in the inner (non-fluorescent) region of the colony, compared with the outer (fluorescent) region (Figure 2E). Alanine conversion into pyruvate and ammonium was also spatially regulated. Two pathways for converting alanine to pyruvate are known: The reversible conversion by enzymes involved in the alanine biosynthetic pathways, and the irreversible conversion mediated by the *dadAX* operon (Figure 2—figure supplement 4B). The latter encodes a racemase (*dadX*) and a dehydrogenase (*dadA*) (Lobocka et al., 1994). In colonies grown for 72 h, the expression of the *dadAX* operon was down-regulated in the inner region, compared with the outer region (Figure 2E). In addition, *dadAX* expression was up-regulated in the whole-colony temporal transcriptomes (6-fold for *dadA* and 5-fold for *dadX*), when comparing 72-h-colonies to 12-h-colonies (Figure 2D). These results indicate that colonies globally up-regulate alanine export and degradation during development and that the anaerobic region of the colony likely exports alanine, while the aerobic region likely converts alanine into pyruvate and ammonium.

### Alanine is exported in anoxic conditions and can be used as a carbon and nitrogen source in oxic conditions

The spatial transcriptomes suggest that alanine is primarily secreted in the anaerobic region of the biofilm. To test this, we explored under which combination of carbon/nitrogen/oxygen availability *E. coli* secretes alanine in liquid conditions. Mass spectrometry measurements from culture supernatants clearly showed that alanine is only secreted under anoxic conditions with glucose and ammonium (Figure 3A). This environment corresponds to the anaerobic base of the colony, where cells are in contact with the glucose- and ammonium-rich M9 agar, suggesting that alanine is secreted in this region, which is consistent with the spatial transcriptome results.

**Figure 3.**
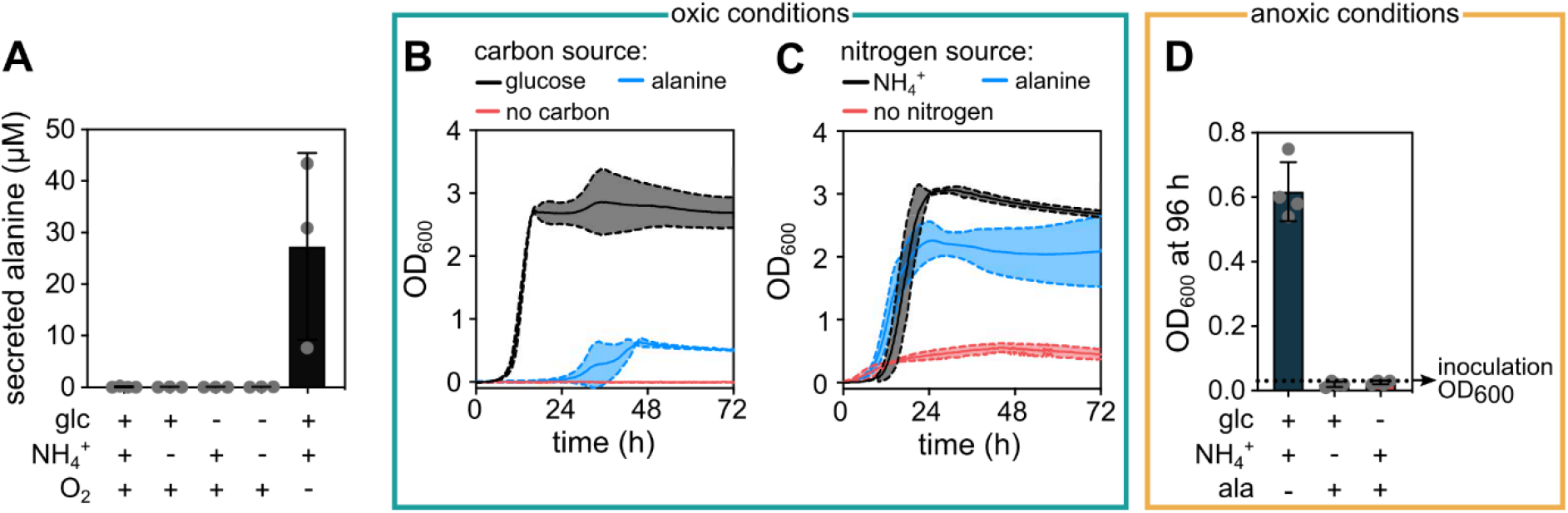
Alanine can be secreted in anoxic conditions and consumed in oxic conditions by *E*. *coli.* (**A**) Extracellular alanine concentration in the supernatant of liquid cultures grown in presence or absence (indicated by + or −) of glucose (glc), ammonium (NH_4_^+^), and molecular oxygen (O_2_). Data are mean ± s.d., *n* = 3. (**B**) Growth curves using M9 minimal salts medium with ammonium as nitrogen source and different carbon sources: Either glucose (5 g/L, as in our standard M9 medium), alanine (10 mM), or no carbon source. (**C**) Growth curves using M9 minimal salts medium with glucose as a carbon source and different nitrogen sources: Either ammonium (22.6 mM, as in our standard M9 medium), alanine (5 mM) or no nitrogen source. For panels **B** and **C**, continuous middle lines correspond to the mean and the dotted lines to the standard deviation, *n* = 3. The growth curves shown for alanine in panels *B* and *C* correspond to the alanine concentrations that resulted in the highest final optical density at 600 nm (OD_600_) in each condition. (**D**) *E. coli* cultures starting with OD_600_ = 0.03 were incubated for 96 h in different anoxic media. The medium contained combinations of glucose, NH_4_^+^, and alanine as indicated. Alanine was used either as the sole nitrogen source (middle bar) or sole carbon source (right bar) with a concentration of 5 mM and 10 mM respectively. The final OD_600_ after 96 h of incubation is shown, corresponding to stationary phase of the culture without alanine. Continuous measurements of OD_600_ in anoxic conditions was not possible in our laboratory due to technical limitations. Data are mean ± s.d., *n* = 4.

The spatial transcriptomes also suggest that alanine is primarily consumed in the aerobic region of the colony. It is well-known that *E. coli* can use extracellular alanine as a carbon or nitrogen source in oxic conditions (Franklin & Venables, 1976; Kornberg, 1965), which we confirmed for our strain and growth conditions by replacing either glucose or ammonium with alanine in the liquid shaking M9 medium (Figure 3B,C). We found that exogenous alanine can be utilized as a poor carbon source, but as a good nitrogen source in oxic conditions. Interestingly, we observed that under anoxic conditions, *E. coli* cannot utilize alanine as a carbon or nitrogen source (Figure 3D). Together, these results suggest that the alanine secreted in the glucose- and ammonium-rich anaerobic region of the colony can be consumed in the aerobic region of the colony as a cross-fed metabolite.

### Bacterial survival in the aerobic region of the colony is influenced by alanine export and consumption

To determine how interference with alanine export and consumption affects colony growth, we created individual and combinatorial deletions of known alanine transport and degradation genes. None of these deletions affected the cellular growth rates in liquid culture, or caused clear phenotypes in colony height or radius after 72 h of incubation on M9 agar (Figure 4—figure supplement 1). However, the mutants displayed substantial differences when we measured the fraction of dead cells in the aerobic region (Figure 4A and Figure 4—figure supplement 2), using a fluorescent nucleic acid stain that can only penetrate the disrupted membranes of dead cells (SYTOX Green). We limited our measurements to the air-facing aerobic region of the colony, because in this region the SYTOX Green fluorescence can be reliably quantified without microscopy artefacts that may occur deeper inside the colony due to poor laser penetration. We observed that the colonies displayed only a low fraction of dead cells at the bottom edges of the colony, but cell death increased towards the top of the colony. Interestingly, the aerobic region at around 50% of the maximum colony height displayed increased cell death when cells carried the Δ*alaE*Δ*dadAX* deletion (Figure 4A), which is a strain that should have a strongly reduced capability for secreting and consuming alanine.

**Figure 4.**
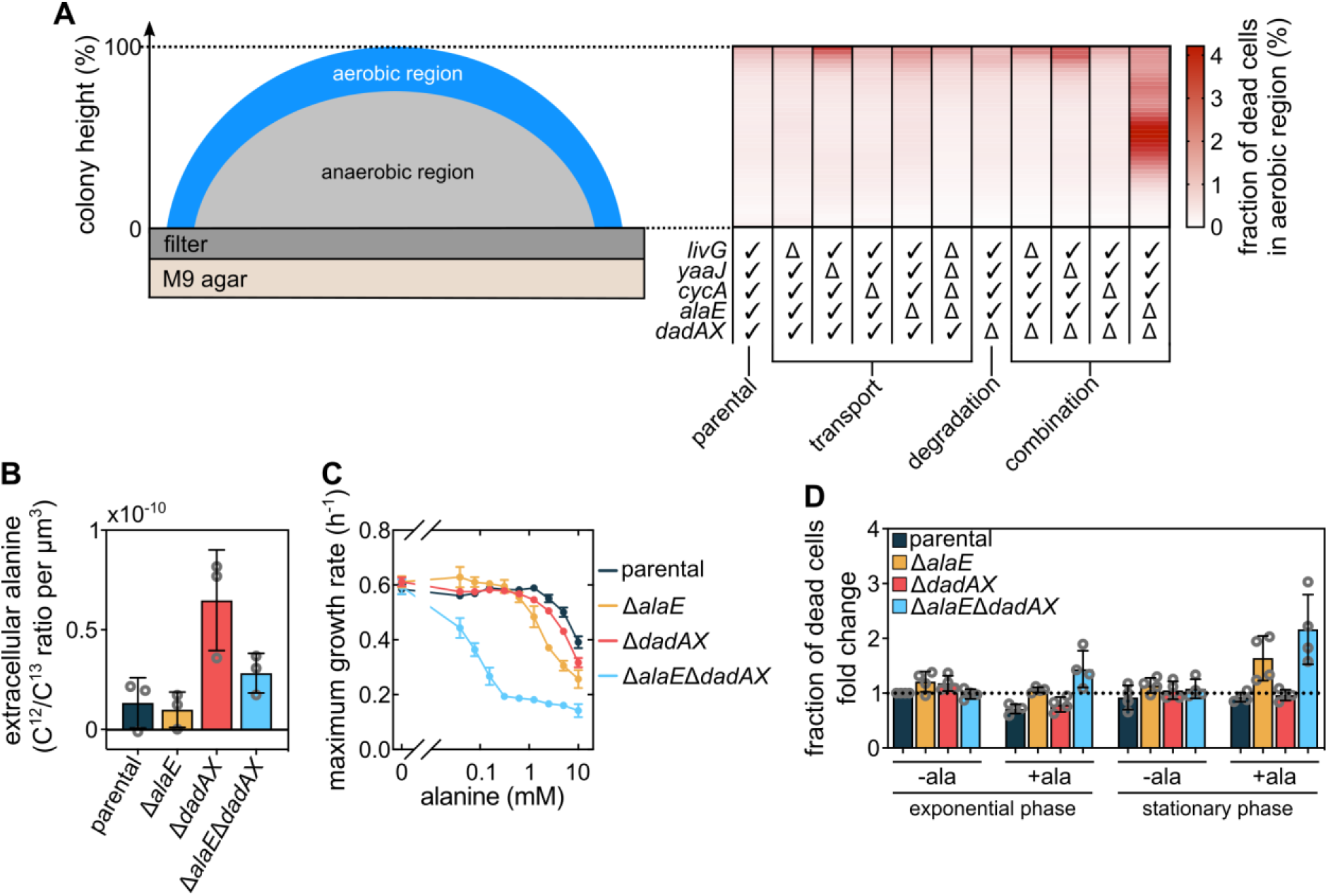
Alanine influences cell viability and growth. (**A**) Measurement of the fraction of dead cells as a function of height in colonies grown for 72 h on M9 agar. These measurements were performed only for cells located within 30 μm from the outer colony surface, which is a conservative measurement of the aerobic region (Figure 2A). The table designates the genotype of strains that were investigated: “✓” indicates that the gene is intact and “Δ” that the gene was deleted. Data in the heatmaps shows means, *n* = 3. Errors are shown in Figure 4—figure supplement 2. (**B**) Extracellular alanine levels measured from colonies (mean ± s.d., *n* = 3). (**C**) Maximum liquid culture growth rate (in presence of glucose and ammonium) for different strains, as a function of the concentration of exogenously added alanine. Data are mean ± s.d., *n* = 3. (**D**) Fraction of dead cells, measured using SYTOX Green fluorescence normalized by OD_600_, in cultures grown with glucose and ammonium in presence or absence of 5 mM alanine. Fold-changes are calculated relative to the parental strain in exponential phase without alanine. Measurements were performed when cultures reached half of the maximum OD_600_ (for exponential phase measurements) or when they reached their maximum OD_600_ (for stationary phase measurements). Bars indicate mean ± s.d., *n* = 4.

To determine if the increased cell death displayed by the Δ*alaE*Δ*dadAX* mutants could be caused by an impaired alanine cross-feeding, we first measured the extracellular alanine concentration in colonies of the relevant mutants using mass spectrometry (Figure 4B). As expected, mutants incapable of the major alanine degradation pathway (Δ*dadAX*) displayed substantially higher extracellular alanine than cells that are impaired for alanine secretion as well (Δ*alaE*Δ*dadAX*). Amino acid transporters can often function as both importers and exporters, yet previous measurements have shown that AlaE acts as an exporter in liquid cultures (Hori, Yoneyama, et al., 2011). The higher extracellular alanine levels of the Δ*dadAX* strain compared with the Δ*alaE*Δ*dadAX* strain (Figure 4B) showed that AlaE also acts as an alanine exporter inside colonies. The parental strain and the Δ*alaE* mutant displayed similarly low extracellular alanine levels, close to the detection limit of our mass spectrometry technique. Interestingly, and consistent with the results of Hori *et al*. in liquid cultures (Hori, Ando, et al., 2011; Hori, Yoneyama, et al., 2011), our detection of extracellular alanine in colonies of the Δ*alaE* and Δ*alaE*Δ*dadAX* mutants indicates that another mechanism for alanine export might exist that is currently unknown.

For cells that lack both the major alanine exporter AlaE and the major alanine degradation pathway *via* DadA and DadX, we hypothesized that the presence of extracellular alanine might lead to an accumulation of intracellular alanine to toxic levels. It has previously been shown that excess levels of intracellular alanine can inhibit growth (Katsube et al., 2019). Indeed, we observed that all strains show a decreased growth rate in liquid media containing high levels of alanine, yet strains carrying the Δ*alaE*Δ*dadAX* mutation were much more sensitive to exogenously added alanine than the parental strain (Figure 4C). This result was not due to unspecific effects of alanine (such as osmolarity changes), because no significant differences between the mutants and the parental strain were observed when serine was added exogenously instead of alanine (Figure 4—figure supplement 3). Therefore, extracellular alanine can modulate bacterial growth rates, particularly for strains that are deficient in alanine cross-feeding (Δ*alaE*Δ*dadAX*).

Since colonies can accumulate alanine in their extracellular space (Figure 4B) and their growth rate can be reduced by extracellular alanine (Figure 4C), we hypothesized that the increased cell death in the aerobic region of the Δ*alaE*Δ*dadAX* colonies (Figure 4A) is due to the accumulation of toxic extracellular alanine levels in this region, which arise from the impaired cross-feeding of this strain. To test this hypothesis, we measured cell viability for the mutants in liquid cultures with and without exogenous alanine during mid-exponential phase and in stationary phase. We found that in the presence of alanine, Δ*alaE*Δ*dadAX* mutants displayed higher cell death than the parental strain, which was particularly strong in stationary phase conditions (Figure 4D). If cells can still export alanine (Δ*dadAX*), cell viability is significantly improved in the presence of extracellular alanine, compared with the Δ*alaE*Δ*dadAX* mutant. If cells can still degrade alanine (Δ*alaE*), cell viability is only slightly improved under stationary phase conditions, compared with the Δ*alaE*Δ*dadAX* mutant. These observations are consistent with degree of sensitivity to extracellular alanine of the Δ*alaE*Δ*dadAX* and Δ*alaE* mutants (Figure 4C). We speculate that the exponentially growing cultures resembled the aerobic periphery at the base of the colony that is in contact with glucose and ammonium, whereas the stationary phase cultures resemble the aerobic region above, which is nutrient depleted if no cross-feeding is present.

Together, these results support the hypothesis that in Δ*alaE*Δ*dadAX* mutant colonies, extracellular alanine accumulates due to impaired cross-feeding, which causes the increased cell death compared with the parental strain. Alanine cross-feeding could therefore have an important effect for the aerobic region above the nutrient-rich base of the colony.

### Alanine cross-feeding influences colony morphology

If the alanine that is secreted in the anaerobic base of the colony is consumed in aerobic higher regions of the colony, alanine could serve as a carbon and nitrogen source and support growth in this region. From measurements of the colony height and diameter of the colony base for mutants impaired in cross-feeding (Figure 4—figure supplement 1), we know that alanine cross-feeding does not contribute to aerobic growth at the very top of the colony or at the outer edge of the base. We therefore investigated effects of alanine cross-feeding on cellular growth in the aerobic region at mid-height, where the fraction of dead cells was highest for the Δ*alaE*Δ*dadAX* mutants (Figure 4A). Measurements of the colony morphology showed that Δ*alaE*Δ*dadAX* and Δ*alaE* mutants displayed a significantly decreased colony curvature in the relevant region, in comparison to the parental strain (Figure 5A,B). The detection of this phenotypic difference in colony morphology was enabled by the high reproducibility of our *E. coli* colony biofilm growth assay. The “bulge” of the parental colonies suggests that this region grows due to alanine cross-feeding. This interpretation is consistent with recent simulations of colony growth without cross-feeding, which resulted in nearly conical colony shapes with triangular *xz*-cross sections (Warren et al., 2019) that lacked the “bulge” morphology we observed for the cross-feeding capable strains.

**Figure 5.**
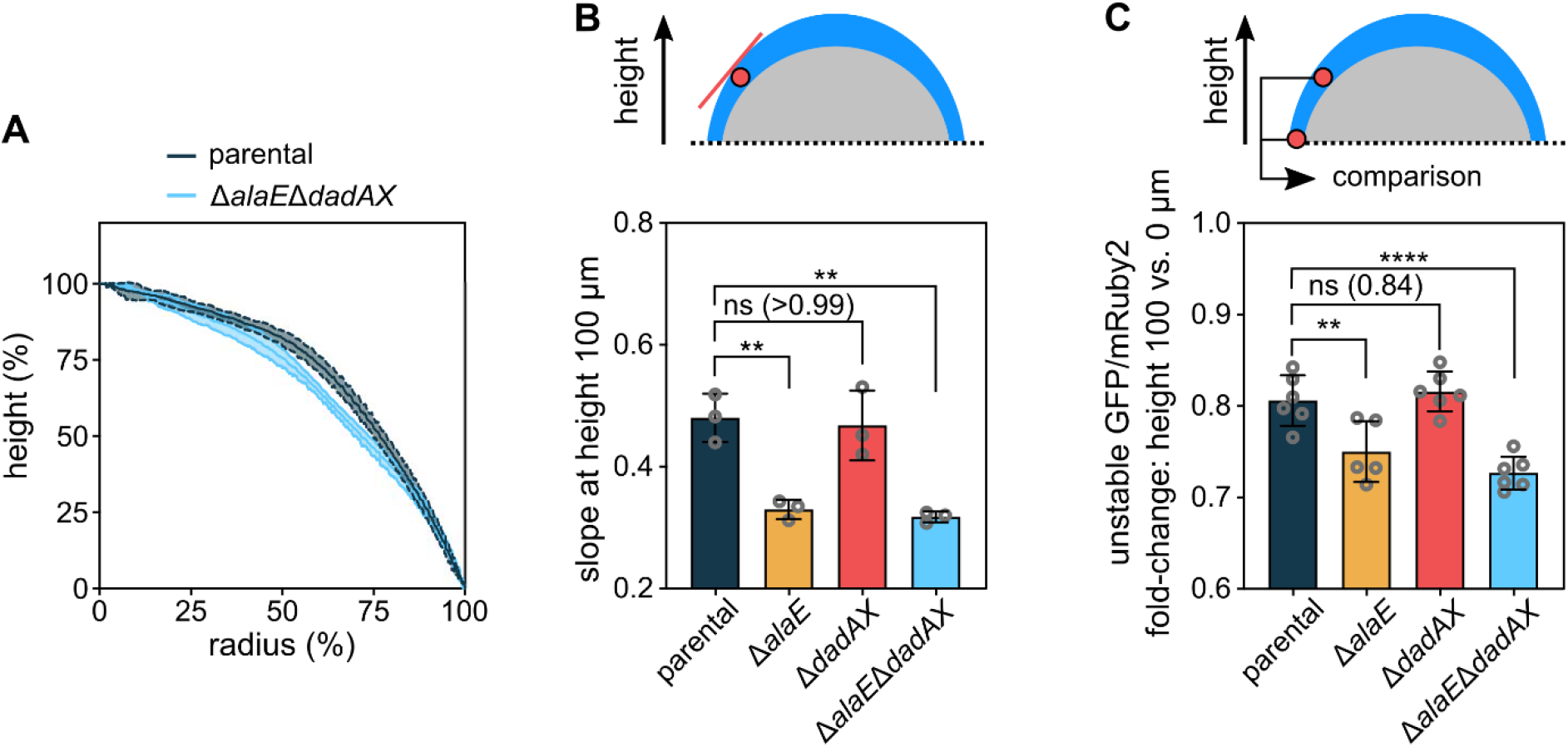
Colony morphology is influenced by alanine cross-feeding. (**A**) Colony height as a function of the colony radius, for colonies grown for 72 h. Central lines correspond to the mean and the shaded area to the s.d., *n* = 3. (**B**) Slope of curves in panel *A* at height 100 μm. Data are mean ± s.d., *n* = 3. (**C**) Fluorescence ratio of constitutively expressed unstable GFP and stable mRuby2, which is a measure for the cellular growth rate, as shown in Figure 5—figure supplement 1. The unstable GFP was constructed by adding the ASV-tag to superfolder GFP. The fold-change of this fluorescent protein ratio was calculated for the aerobic colony region between the heights 100 μm and 0 μm. The fluorescent protein ratio was only measured for cells located within 30 μm from the outer colony surface, as a conservative measure of the aerobic region. Data are mean ± s.d., *n* = 5-6. Statistical significances were calculated using one-way ANOVA with Dunnett’s correction. Non-significant differences are labelled “ns” and the *p*-value is shown in brackets. \*\**p* < 0.01, \*\*\*\**p* < 0.0001.

To further experimentally investigate the effect of alanine cross-feeding on cellular growth in the mid-height aerobic region, we measured the fluorescence ratio of an unstable version of the superfolder GFP (Andersen et al., 1998) and the long-lived mRuby2, both of which were expressed constitutively using a P_*tac*_ promoter. The ratio of these fluorescent proteins serves as a measure of the cellular growth rate (Figure 5—figure supplement 1). These measurements showed that the parental strain grows faster in the mid-height aerobic region than the cross-feeding impaired Δ*alaE*Δ*dadAX* mutant (Figure 5C), which further supports the hypothesis that this region of the colony relies on alanine as a carbon and nitrogen source.

## Discussion

We showed that alanine metabolism is spatiotemporally regulated inside *E. coli* colony biofilms. The high degree of control and reproducibility of our model system enabled us to determine that interference with the export and consumption of alanine causes phenotypes in cell viability and cell growth rate. These viability and growth rate phenotypes are localized in the region of the colony that depends on alanine as a nutrient source, but they affect the global morphology of the colony.

Based on our results, we propose the following model for the spatial organization of alanine metabolism in colonies that have grown for 72 h (Figure 6): Cells at the bottom periphery of the colony (red region in Figure 6) have access to oxygen, glucose and ammonium, and perform aerobic respiration (Basan et al., 2015; Cole et al., 2015). Cells at the bottom center of the colony (orange region in Figure 6) are anaerobic yet they have access to glucose and ammonium from the agar-solidified medium. These cells ferment glucose and secrete alanine, primarily *via* AlaE. Although many amino acid exporters have been described for *E. coli* (39, 42, 43), their functions have remained elusive under regular physiological conditions. Our data now reveal a function for the alanine exporter AlaE during biofilm growth. Secreted alanine diffuses through the colony and can only be utilized by aerobic nutrient-deficient cells (blue region in Figure 6). Alanine consumption in the mid-height aerobic region also has a detoxification effect, by reducing otherwise inhibitory levels of extracellular alanine. Alanine consumption at the aerobic top of the colony is not significant, perhaps because the extracellular alanine is consumed before it reaches this region.

**Figure 6.**
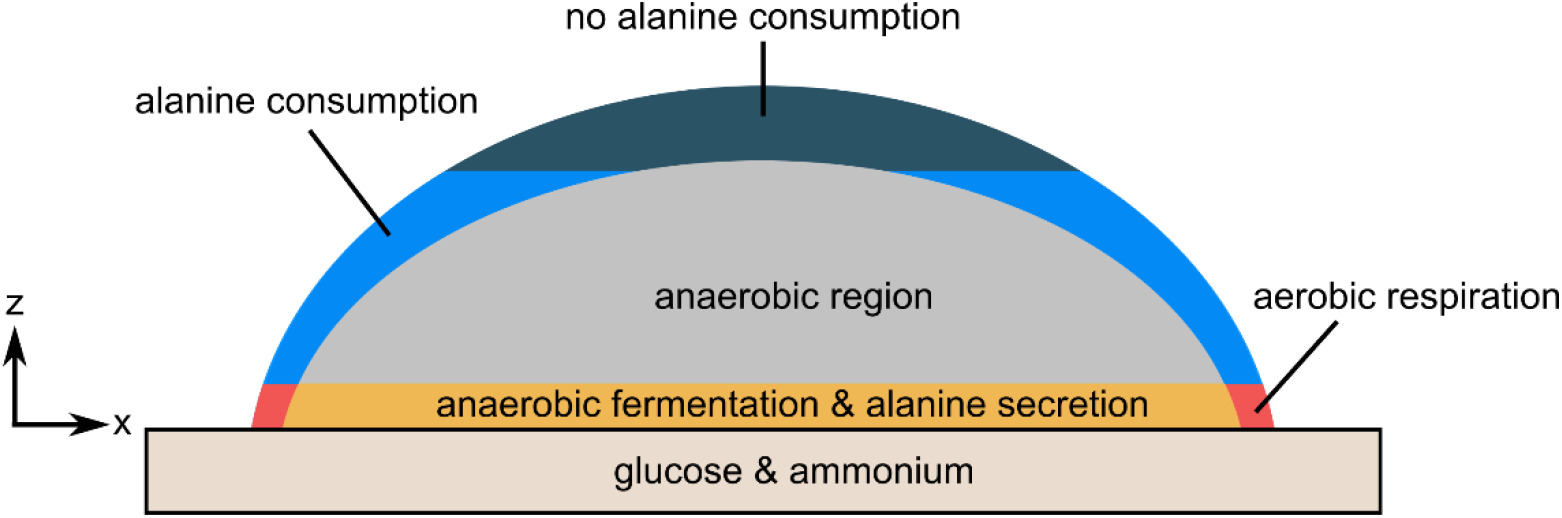
Model for alanine cross-feeding in *E. coli* colony biofilms. This model applies to colonies grown for 72 h on solid M9 agar containing glucose and ammonium. Cells in the bottom layer of the biofilm have access to glucose and ammonium. Only cells in the outer periphery of the biofilm have access to oxygen. Cells in the red region can use ammonium, glucose, and oxygen to perform aerobic respiration. Cells in the orange region have access to glucose and ammonium, but no oxygen. These cells secrete alanine. The secreted alanine can be consumed by the aerobic cells above this layer (depicted as blue), which convert alanine into pyruvate and ammonium that can be used for growth and to maintain cell viability.

The spatiotemporal organization of alanine metabolism during colony growth is unique among amino acids (Figure 1D,E), and is based on the secretion of alanine in the anaerobic, glucose-rich, and ammonium-rich base of the colony. Why is alanine secreted in this region? We speculate that this is not an altruistic trait evolved to support a starving aerobic subpopulation at a different location, because such a trait would be highly susceptible to social cheaters in a multi-species community. Instead, alanine secretion in this region may be a necessity to avoid high intracellular alanine levels that presumably result from anaerobic fermentation of glucose in the presence of ammonium. High alanine levels are inhibitory (Figure 4C), so that secretion of alanine and transport of alanine away from this population is beneficial to this population.

On the other side of the cross-feeding interaction, the alanine-consuming subpopulation in the aerobic mid-height region of the colony strongly benefits from the secreted alanine originating from the base of the colony. We note that the alanine consumption in the aerobic mid-height region of the colony necessarily causes a steeper alanine concentration gradient between the two interacting subpopulations, compared with a case in which no alanine is consumed. Due to Fick’s law of diffusion, a steeper concentration gradient causes a higher diffusive flux. Therefore, the presence of the alanine-consuming subpopulation results in a benefit for the alanine-secreting subpopulation, by causing a higher diffusive flux of alanine away from the alanine-secreting population. It is unclear whether this benefit for the alanine-secreting subpopulation is significant, as it is not possible for us to measure local growth rates or cell viability in the anaerobic base of the colony due to optical limitations of confocal fluorescence microscopy in this region. In summary, the interaction between the two cross-feeding subpopulations is likely mutualistic.

The spatial organization of alanine cross-feeding between two subpopulations we described in this study is analogous to the carbon cross-feeding in *E. coli* colonies *via* acetate, because it involves metabolite secretion in the anaerobic population and consumption in the carbon-starved aerobic population (Cole et al., 2015; Dal Co et al., 2019; Wolfsberg et al., 2018). However, the location of the population that is proposed to consume acetate as a carbon source spans most of the aerobic region of the colony (Cole et al., 2015), whereas alanine is primarily consumed only in the aerobic mid-height region of the colony. Interestingly, our spatial transcriptomes did not reveal a signature for acetate cross-feeding between the anaerobic and aerobic regions of the colony (Figure 1—figure supplement 4B), yet transcripts coding for enzymes involved in lactate, formate, and succinate metabolism display patterns that are indicative of spatially organized metabolism that could be the basis of carbon-cross-feeding. In contrast to these metabolites, alanine can not only be used as carbon source, but also as a nitrogen source in the cross-feeding-dependent region, and we note that it is currently not clear what limits growth in the higher regions of the colony – whether it is carbon, nitrogen, or other elements such as iron, sulfur, or phosphorous. Very recently, it was shown that alanine can be cross-fed in colonies of the Gram-positive bacterium *Bacillus subtilis*, and that this effect required three-dimensional colonies for unknown reasons (Arjes et al., 2021). In our *E. coli* model system, three-dimensional growth is required to create an anaerobic region replete with carbon and nitrogen sources that causes alanine secretion, and we speculate that this effect may also be required in *B. subtilis,* and in many other species.

Several cross-feeding interactions have been described in detail for multi-species communities (Henson et al., 2019; Kim et al., 2017; Moree et al., 2012; Pande et al., 2016; Røder et al., 2020; Smith et al., 2019; Watrous et al., 2013; Yang et al., 2009). Even though cross-feeding interactions are likely a ubiquitous process in single-species bacterial multicellular structures (San Roman & Wagner, 2018), only a few metabolic interactions between subpopulations have been documented for single-species biofilms (Arjes et al., 2021; Cole et al., 2015; Evans et al., 2020; Lin et al., 2018; Liu et al., 2015). For *E. coli* colonies grown on agar-solidified medium, and for bacterial communities in general, it is still unclear how many subpopulations interact metabolically, and on which length and time scales these interactions take place, and how significant these many interactions are for the community growth and stability.

## Conclusion

In this study, we employed an unbiased approach based on temporal and spatial transcriptomes and metabolomes to reveal that a multitude of amino acids and mixed acid fermentation pathways display profiles that are consistent with cross-feeding. Particularly strong regulation was displayed by alanine metabolism, and we showed that alanine is a cross-fed metabolite inside *E. coli* colonies, between two spatially segregated subpopulations, with an interaction length scale of tens of microns. Alanine consumption supports growth in the cross-feeding-dependent region of the colony as a carbon and nitrogen source. Although many aspects of metabolism in biofilms are still unknown, methods for improved spatial and temporal analyses of metabolite profiles and transcriptome data promise the possibility to discover new metabolic interactions, and more generally understand the stability and functions of microbial communities.

## Acknowledgements

We are grateful to Thorben Schramm and Nicole Paczia for performing additional metabolite measurements. We thank Gabriele Malengo for helping with the cell sorting procedure. We also thank Miriam Bayer, Praveen K. Singh, Daniel Rode, Lucia Vidakovic, Sanika Vaidya, and all members of the Drescher lab for their comments and fruitful discussions. This work was supported by grants from the European Research Council (StG-716734), Deutsche Forschungsgemeinschaft (SFB987 and DR 982/5-1), Minna James Heineman Foundation, Bundesministerium für Bildung und Forschung (TARGET-Biofilms), Max Planck Society, and the Swiss National Science Foundation NCCR “AntiResist” (to K.D.), and an NSF CAREER award and NIH/NIAID grant R01AI103369 (to L.E.P.D.).

## Authors Contributions

F.D.-P. and K.D. designed experiments and analyzed data. F.D.-P. performed growth and microscopy-based experiments. F.D-P., M.L., and H.L. designed and performed metabolomic experiments. Ka.No. designed and performed transcriptome experiments. H.J., R.H. and E.J. developed and performed image analysis. J.K.J. and L.E.P.D. designed and performed oxygen measurements. Ko.Ne. and R.H. developed microscope control and imaging techniques. M.F.H. generated strains. F.D.-P., and K.D. created the figures. A.P-W., L.E.P.D., and K.D. wrote the manuscript with input from all authors. All authors made important conceptual contributions to the project and the interpretation of results, often in group discussions. K.D. supervised and coordinated the project.

## Materials and Methods

### Strains, strain construction, and media

All *E. coli* strains used in this study are derivatives of the *E. coli* K-12 AR3110 strain (Serra, Richter, & Hengge, 2013). For the construction of plasmids and bacterial strains, standard molecular biology techniques were applied (Sambrook et al., 1989), using enzymes purchased from New England Biolabs or Takara Bio. All AR3110 derivatives carried a constitutively expressed fluorescent protein expression system (based on the P_*tac*_ promoter without the *lac* operator) inserted in the chromosome at the *attB* site. To generate chromosomal deletions, the lambda red system was used to replace the target region with an antibiotic cassette flanked by FRT sites. Then, the Flp-FRT recombination system was utilized to remove the antibiotic resistance cassette. To insert the unstable fluorescent protein sfGFP(ASV) (Andersen et al., 1998) expressed under the control of the P_*tac*_ promoter (without the *lac* operator) and a chloramphenicol cassette flanked by FRT sites into the Tn7 insertion site, we used the protocol described by Choi *et.al.* (Choi et al., 2005). After the fragment was inserted, the chloramphenicol resistance cassette was removed using the Flp-FRT recombination system. All strains, plasmids, and oligonucleotides that were used in this study are listed in Table S1, Table S2, and Table S3, respectively.

Cultures were grown in LB-Miller medium (10 g L^−1^ NaCl, 10 g L^−1^ tryptone, 5 g L^−1^ yeast extract) for routine culture and cloning, or in M9 minimal salts medium supplemented with 5 g L^−1^ D-glucose (which we refer to as “M9 medium” throughout the article for simplicity). The M9 minimal salts medium consisted of the following components: 42.3 mM Na_2_HPO_4_ (Carl Roth P030.2), 22 mM KH_2_PO_4_ (Carl Roth 3904.1), 11.3 mM (NH_4_)_2_SO_4_ (Carl Roth 3746.2), 8.56 mM NaCl (Carl Roth HN00.2), 0.1 mM CaCl_2_ (Sigma-Aldrich C5670), 1 mM MgSO_4_, (Sigma-Aldrich M2643), 60 μM FeCl_3_ (Sigma-Aldrich 31332), 2.8 μM thiamine-HCl (Sigma-Aldrich T4625), 6.3 μM ZnSO_4_ (Sigma-Aldrich Z0251), 7 μM CuCl_2_ (Sigma-Aldrich 307483), 7.1 μM MnSO_4_ (Sigma-Aldrich M8179), 7.6 μM CoCl_2_ (Carl Roth 7095.1). M9 agar plates were made using 8 mL of M9 medium (as defined above) with 1.5 % w/v agar-agar, aliquoted into petri dishes with 35 mm diameter and 10 mm height (Sarstedt 82.1135.500).

### Colony biofilm growth

Cultures were inoculated from frozen glycerol stocks kept at −80 °C into LB-Miller medium with kanamycin (50 μg mL^−1^) and grown for 5 h at 37 °C with shaking at 220 rpm. At this point, 1 μL of the culture was inoculated into 5 mL of M9 medium inside a 100-mL-Erlenmeyer flask and grown at 37 °C with shaking at 220 rpm for 16-22 h. The cultures were continuously kept in exponential phase by regular back-dilutions, with optical density at 600 nm (OD_600_) always below 0.6. Aliquots from these cultures were passed through a sterile 0.45 μm pore size polyvinylidene fluoride membrane filter of diameter 5 mm, unless stated otherwise. High-resolution confocal fluorescence microscopy showed that this treatment resulted in spatially well-separated single cells on the filter membrane. Using clean and sterile stainless-steel tweezers, these filter membranes carrying the cells were immediately placed directly onto M9 agar plates and incubated at 37 °C for up to 72 h.

### Microscopy

All imaging was performed using a Yokogawa spinning disk confocal unit mounted on a Nikon Ti-E inverted microscope. A Nikon 40x air extra-long working distance objective with numerical aperture (NA) 0.60 was used for all imaging, except for mutant screening (Figure 4—figure supplement where a 4x air objective (NA 0.13, Nikon) was used, and single-cell imaging (Figure 2—figure supplement 1A) were a 100x oil objective (NA 1.45, Nikon) was used. All imaging was done inside a microscope incubator kept at 37 °C. Instead of imaging through the filter, colonies were imaged facing down and the Petri dish lid was removed (Figure 1—figure supplement 1A). To maintain a high humidity and avoid evaporation of the M9 agar, the space between the microscope objective and the petri dish was sealed with flexible plastic foil. To avoid condensation on the objective, the objective was heated using an objective heater at 37 °C.

### Colony detection and biovolume measurements with adaptive microscopy

In preliminary experiments, we observed that colony growth of the wild type *E. coli* strain AR3110 resulted in heterogeneous shapes (Figure 1—figure supplement 1B) that were caused by flagella-based motility of cells on the filter membranes in the very early stages of incubation. To avoid the effects caused by cellular motility during early colony growth, all subsequently experiments were performed using a strain that lacks the flagellin FliC (Δ*fliC*), which we used as parental strain in this study. This strain was incapable of swimming motility, and the resulting colonies were highly reproducible in shape and highly symmetric (Figure 1B and Figure 1—figure supplement 1C).

To determine the biomass of colonies, the colonies were grown on filter membranes on M9 agar as explained above, but with the addition of 0.2 μm dark red fluorescent beads (Invitrogen, F8807) to the bacterial suspension prior to filtering. This resulted in far-red fluorescent beads being located on the filter in addition to the red fluorescent cells. Using adaptive microscopy (Jeckel & Drescher, 2020) these beads were used to find the correct focal plane for imaging of the bacterial cells (or base of the colonies). After 12 h, 18 h, 24 h, 32 h, 42 h, 48 h, 60 h, and 72 h of colony growth, the colonies were imaged in 3D using confocal microscopy. All colonies on the membrane filter were imaged using an adaptive microscopy approach (Jeckel & Drescher, 2020) as follows: The whole filter was scanned using 2D imaging, followed by an identification of all colonies on the filter, followed by a high-resolution 3D imaging of each colony. From the scans of the whole filter, the number of colonies was determined. From the 3D images of the colonies, the biovolume was calculated. To obtain the biovolume, the outline of the colony was identified by thresholding the image gradient in each *z*-slice. The convex area of this binary image was then used as a measure for the biomass present in this plane such that summation over all slices followed by multiplication with the appropriate μm^3^/voxel calibration yielded the biovolume of each colony.

### Liquid growth assays

Cultures were inoculated from −80 °C frozen stocks into LB-Miller media and grown for 5 h at 37 °C with shaking at 220 rpm. Each culture was back-diluted 5,000-fold into 5 mL of M9 medium inside a 100-mL-Erlenmeyer flask, and grown at 37 °C with shaking at 220 rpm. At an OD_600_ of 0.3 each culture was washed 3 times in M9 minimal salts medium (lacking glucose and ammonium sulfate) and resuspended in the same volume of the medium of interest. These bacterial suspensions were diluted 10-fold and transferred into a 96-well plate (Sarstedt, 82.1581.001), and incubated at 37 °C with shaking in a microtiter plate reader (Epoch2, Biotek). The resulting growth curve data was analysed using Matlab (version R2019b, Mathworks).

To simultaneously measure OD_600_ and the ratio between unstable GFP (with the ASV-tag) and mRuby2 as a function of time (Figure 5—figure supplement 1), strain KDE2937 was inoculated from a −80 °C frozen stock into LB-Miller media and grown for 5 h at 37 °C with shaking at 220 rpm. The culture was then back-diluted 5,000-fold into 5 mL of M9 medium inside a 100-mL Erlenmeyer flask and grown for 16 h. This culture was used to inoculate 75 mL of M9 medium inside a 1-L Erlenmeyer flask with an adjusted OD_600_ of 0.05. This culture was grown at 37°C with shaking at 220 rpm. Aliquots of the culture were taken every 30 min, to measure the OD_600_ and to determine the ratio between unstable GFP (with the ASV-tag) and mRuby2 using microscopy. To image the aliquots, they were placed between a cover slip and a M9 agar pad and imaged as described in the microscopy methods section.

### Anoxic liquid growth assays

Cultures were inoculated from −80 °C frozen stocks into LB-Miller media and grown for 5 h at 37 °C with shaking at 220 rpm. Each culture was then back-diluted 5,000-fold into 5 mL of M9 medium inside a 100-mL Erlenmeyer flask, and grown at 37 °C with shaking at 220 rpm. At OD_600_ = 0.7 each culture was washed 3 times in M9 minimal salts medium (lacking glucose and ammonium sulfate) and resuspended in 300 μL. These bacterial suspensions were used to inoculate 20 mL of the particular medium under investigation with a starting OD_600_ of 0.03, which was placed into a 50-mL bottle that was closed with a gas-tight lid containing a rubber septum (Duran, 292062803). Immediately after inoculation, the bottle and the culture were made anoxic using the following protocol: A needle connected to a system that can switch between applying a vacuum and gaseous N_2_ was introduced through the rubber septum. Then, vacuum was applied to remove the gas in the bottle for 1 min, followed by flushing the bottle with N_2_ at 1 bar for 1 min, while the medium in the bottle was continuously mixed using a magnetic stirrer. This process was repeated for 15 cycles. After this process, the needle was slowly removed from the bottle and the cultures were incubated at 37 °C with shaking at 220 rpm. After 96 h of incubation, the OD_600_ was measured (Figure 3D).

### Measurement of extracellular alanine in liquid culture supernatants

Cultures were inoculated from frozen −80 °C stocks into LB-Miller media with kanamycin (50 μg mL^−1^) and grown for 5 h at 37 °C with shaking at 220 rpm. The cultures were then back-diluted 5,000-fold into 5 mL of M9 medium inside a 100-mL-Erlenmeyer flask and incubated at 37 °C with shaking at 220 rpm. These cultures were kept in exponential phase by regular back-dilutions, with optical density at 600 nm (OD_600_) always below 0.6. At an OD_600_ of 0.3 the cultures were centrifuged at 14,000 g for 2 min and the supernatant was removed. To investigate aerobic growth conditions, the pellets were suspended in M9 minimal salts medium supplemented with glucose and the OD_600_ was adjusted to 0.1 by dilution. For aerobic starvation conditions, the pellets were suspended in same volume of M9 medium lacking ammonium sulfate but including glucose, M9 medium lacking glucose but including ammonium sulfate, or M9 medium lacking both glucose and ammonium sulfate. The cultures were then placed into 100-mL-Erlenmeyer flasks at 37 °C with shaking at 220 rpm as before. To investigate anaerobic conditions, the pellets were suspended in the same media as described above, but the resulting cultures were transferred into a closed 15-mL-conical centrifuge tube (Sarstedt, 62.554.100) filled to the top. The tubes were oriented horizontally and incubated at 37 °C with shaking at 100 rpm. For the aerobic and anaerobic starvation conditions, the samples were taken after 2 h of incubation. For aerobic and anaerobic conditions that permitted growth, samples were taken during exponential growth phase. Samples were processed and analysed using mass spectrometry as described below.

### Sample processing for mass spectrometry-based metabolomics

To measure metabolites in whole colonies over time, filter membranes that carried colonies were transferred into 150 μL of a mixture of 40:40:20 (v/v) acetonitrile:methanol:water at −20 °C for metabolite extraction. This suspension was vortexed with a glass bead to disrupt the colonies.

To measure extracellular metabolites from colonies, the filter membranes carrying the colonies were resuspended in 1 mL phosphate-buffered saline (PBS; 8 g l^−1^ NaCl, 0.2 g ^l−1^ KCl, 1.44 g l^−1^ Na_2_HPO_4_, 0.24 g l^−1^ KH_2_PO_4_, pH 7.4) at 37 °C. In this case, no glass bead vortexing was needed, as the colonies readily dissolved. The suspension was immediately vacuum-filtered using a 0.45 μm pore size filter (HVLP02500, Merck Millipore) and 100 μL of the flow-through were mixed with 400 μL of a mixture of 50:50 (v/v) acetonitrile:methanol at −20 °C.

To measure extracellular metabolites from liquid cultures, 1 mL of grown cultures were filtered on a 0.45 μm pore size filter (HVLP02500, Merck Millipore) and 100 μL of the flow through were mixed with 400 μL of a mixture of 50:50 (v/v) acetonitrile:methanol at −20 °C.

All extracts were centrifuged for 15 minutes at 11,000 g at −9 °C and stored at −80 °C until mass spectrometry analysis.

### Mass spectrometry measurements

For metabolomics, centrifuged extracts were mixed with ^13^C-labeled internal standards. Chromatographic separation was performed on an Agilent 1290 Infinity II LC System (Agilent Technologies) equipped with an Acquity UPLC BEH Amide column (2.1 × 30 mm, particle size 1.8 μm, Waters) for acidic conditions and an iHilic-Fusion (P) HPLC column (2.1 × 50 mm, particle size 5 μm, Hilicon) for basic conditions. The following binary gradients with a flow rate of 400 μl min^−1^ were applied. Acidic condition: 0-1.3 min isocratic 10% A (water with 0.1% v/v formic acid, 10 mM ammonium formate), 90% B (acetonitrile with 0.1% v/v formic acid,),1.3-1.5 min linear from 90% to 40% B; 1.5-1.7 min linear from 40% to 90% B, 1.7-2 min isocratic 90% B. Basic condition: 0-1.3 min isocratic 10% A (water with formic acid 0.2% (v/v), 10 mM ammonium carbonate), 90% B (acetonitrile); 1.3-1.5 min linear from 90% to 40% B; 1.5-1.7 min linear from 40% to 90% B, 1.7-2 min isocratic 90% B. The injection volume was 3.0 μl (full loop injection).

Ions were detected using an Agilent 6495 triple quadrupole mass spectrometer (Agilent Technologies) equipped with an Agilent Jet Stream electrospray ion source in positive and negative ion mode. The source gas temperature was set to 200 °C, with 14 L min^−1^ drying gas and a nebulizer pressure of 24 psi. Sheath gas temperature was set to 300 °C and the flow to 11 L min^−1^. Electrospray nozzle and capillary voltages were set to 500 and 2500 V, respectively. Metabolites were identified by multiple reaction monitoring (MRM). MRM parameters were optimized and validated with authentic standards (Guder et al., 2017). Metabolites were measured in ^12^C and ^13^C isoforms, and the data was analysed with a published Matlab code (Guder et al., 2017).

### Data analysis for metabolomics

For whole-colony metabolite measurements, the mass spectrometry measurements were normalized by the total biovolume measured with confocal microscopy. For measurements of the extracellular metabolites in colonies (Figure 4B), the mass spectrometry measurements were normalized by colony number and average colony volume determined by confocal microscopy. To create heatmaps of the metabolite dynamics, the Genesis software (Sturn et al., 2002) was used.

### Whole-colony transcriptomes

Colonies were grown on filter membranes as described above. After 12 h, 18 h, 24 h, 32 h, 42 h, 48 h, 60 h, and 72 h of growth, filters carrying the colonies were picked up using forceps and transferred into 1.5 mL Eppendorf tubes, which were immediately placed into liquid nitrogen, followed by storage at −80 °C until further processing. To extract the RNA, a glass bead and 600 μL of cell lysis buffer were added during thawing of the sample at room temperature. Cell lysis buffer consisted of TE (10 mM Tris, adjusted to pH 8.0 with HCl, 1 mM EDTA) and 1 mg/mL of chicken egg lysozyme (Sigma, L6876). The colonies were disrupted by vortexing and the cell suspension was moved to a new 1.5 mL Eppendorf tube. Then, total RNA was extracted using the hot SDS/phenol method (Jahn et al., 2008) with some modifications as follows. Cells were lysed at 65 °C for 2 min in the presence of 1% (w/v) SDS, and the lysate was incubated with 750 μL of Roti-Aqua-Phenol (Carl Roth, A980) at 65 °C for 8 min, followed by the addition of 750 μL chloroform (Sigma, C2432) to the aqueous phase and centrifugation using a phase lock gel tube (VWR, 733-2478). RNA was purified from this suspension by ethanol precipitation and dissolved in 60 μL of RNase-free water. Samples were then treated with TURBO DNase (Thermo Fisher, AM2238) and rRNA depletion was performed using Ribo-Zero rRNA Removal Kit for bacteria (Illumina, MRZB12424). Sequencing library preparation was carried out using NEBNext Ultra II Directional RNA Library Prep with Sample Purification Beads (NEB, E7765S). Sequencing was carried out at the Max Planck Genome Centre (Cologne, Germany) using an Illumina HiSeq3000 with 150 bp single reads. All transcriptomic analyses were performed using the software CLC Genomics Workbench v11.0 (Qiagen). The *E. coli* K-12 W3310 genome (Hayashi et al., 2006), parental strain of *E. coli* AR3110 (Serra, Richter, & Hengge, 2013), was used as reference for annotation. For creating heatmaps, clustering of transcripts was performed using the software Genesis (Sturn et al., 2002).

### Spatial transcriptomes

Filter membranes carrying colonies that were grown for 72 h were transferred into 2 mL Eppendorf tubes (one filter per tube). Cells from the colonies on a filter were then quickly suspended in 1 mL PBS by vortexing and pipetting, followed by removal of the filter from the tube, and fixation of the cells by adding formaldehyde (Sigma, F8775) to a final concentration of 4 %, for 10 min at room temperature. Formaldehyde fixation did not affect the transcriptomic profile or the fluorescence intensity (Figure 2—figure supplement 2). Then, the fixed cells were washed three times with PBS and the cell suspension was filtered with a 5 μm pore size filter (Sartorius, 17594) to remove aggregates. Cells were separated using fluorescence-activated cell sorting (BD FACSAria Fusion), using the mRuby2 fluorescence as a signal. After sorting, approximately 10^6^ cells were collected in 10 mL of PBS for each bin. To concentrate the samples, they were vacuum-filtered using a 0.45 μm membrane filter (Millipore, HVLP02500). The filters containing the cells were suspended in 400 μL of the cell lysis buffer (same composition as above). The suspension was frozen in liquid nitrogen and stored at −80 °C until RNA extraction.

Total RNA was extracted using the same protocol as described above (based on the hot SDS/phenol method), but with the following modifications to minimize the number of steps and loss of RNA: after treatment with phenol, the aqueous and organic phases were both directly transferred to a phase lock gel tube (VWR, 733-2478) without centrifugation. Chloroform was added and centrifuged at 15,000 rpm for 15 min at 12 °C. After centrifugation, RNA in the aqueous phase was purified and collected in 10 μL of RNase-free water using Agencourt RNAClean XP (Beckman Coulter, A63987) according to the manufacturer’s recommendations. Samples were then treated with TURBO DNase (Thermo Fisher, AM2238) and rRNA depletion was performed using Ribo-Zero rRNA Removal Kit for bacteria (Illumina, MRZB12424). Library preparation was carried out using NEBNext Ultra II Directional RNA Library Prep with Sample Purification Beads (NEB, E7765S). Sequencing was carried out at the Max Planck Genome Centre (Cologne, Germany) using an Illumina HiSeq3000 with 150 bp single reads.

### Measurements of the fraction of dead cells

To measure the fraction of dead cells within the aerobic region of the colonies, the colonies were grown on filter membranes on M9 agar as described above, but with the exception that SYTOX Green (Thermofisher, S7020) with a final concentration of 2.5 μM was added to the M9 agar plates. SYTOX Green is a nucleic acid stain that can only diffuse into dead cells with a compromised cell membrane. After incubation for 72 h, the mRuby2 fluorescence and the SYTOX Green fluorescence in the colonies was imaged using confocal microscopy. Inside the colony, 3D regions that were fluorescent in both the mRuby2 and SYTOX Green channels were identified *via* thresholding. The fraction of dead cells was calculated as the thresholded volume present in both the mRuby2 and SYTOX Green channels, divided by the thresholded volume of the mRuby2 channel. For the quantification of the fraction of dead cells in the aerobic region of the colony, only cells located within 30 μm from the outer surface of the colony were considered, because only in this region oxygen penetration was high enough to generate a sufficiently strong mRuby2 signal.

To measure the fraction of dead cells in liquid cultures, cultures were grown in M9 medium, which was supplemented with SYTOX Green (2.5 μM), incubated at 37 °C inside 96-well plates with continuous shaking. The OD_600_ and SYTOX Green fluorescence were measured in a microtiter plater reader (Spark 10M, Tecan). The fraction of dead cells was calculated as the SYTOX Green fluorescence normalized by the OD_600_.

### Oxygen measurements

Oxygen concentrations were measured in 72-h-old colonies (grown on M9 agar as described above) using a 25-μm-tip oxygen microsensor (Unisense OX-25) according to the manufacturer’s instructions. Briefly, the oxygen microsensor was calibrated using a two-point calibration. The microsensor was first calibrated to atmospheric oxygen using a calibration chamber (Unisense CAL300) containing water continuously bubbled with air. The microsensor was then calibrated to a “zero” oxygen point using an anoxic solution of 0.1 M sodium ascorbate and 0.1 M sodium hydroxide in water. Oxygen measurements were then taken through the top ≥100 μm of the colony biofilm in 5-μm-steps using a measurement time of 3 s at each position, and a wait time between measurements of 5 s. A micromanipulator (Unisense MM33) was used to move the microsensor within the colony. Profiles were recorded using a multimeter (Unisense) and the SensorTrace Profiling software (Unisense).

### Statistical analysis

Statistical analysis was carried out using GraphPad Prism v8 (GraphPad Software), except for the statistical analysis for transcriptomic data, which was performed using the software CLC Genomics Workbench 11.0 (Qiagen).

## Supplementary Information

**Figure 1—figure supplement 1.**
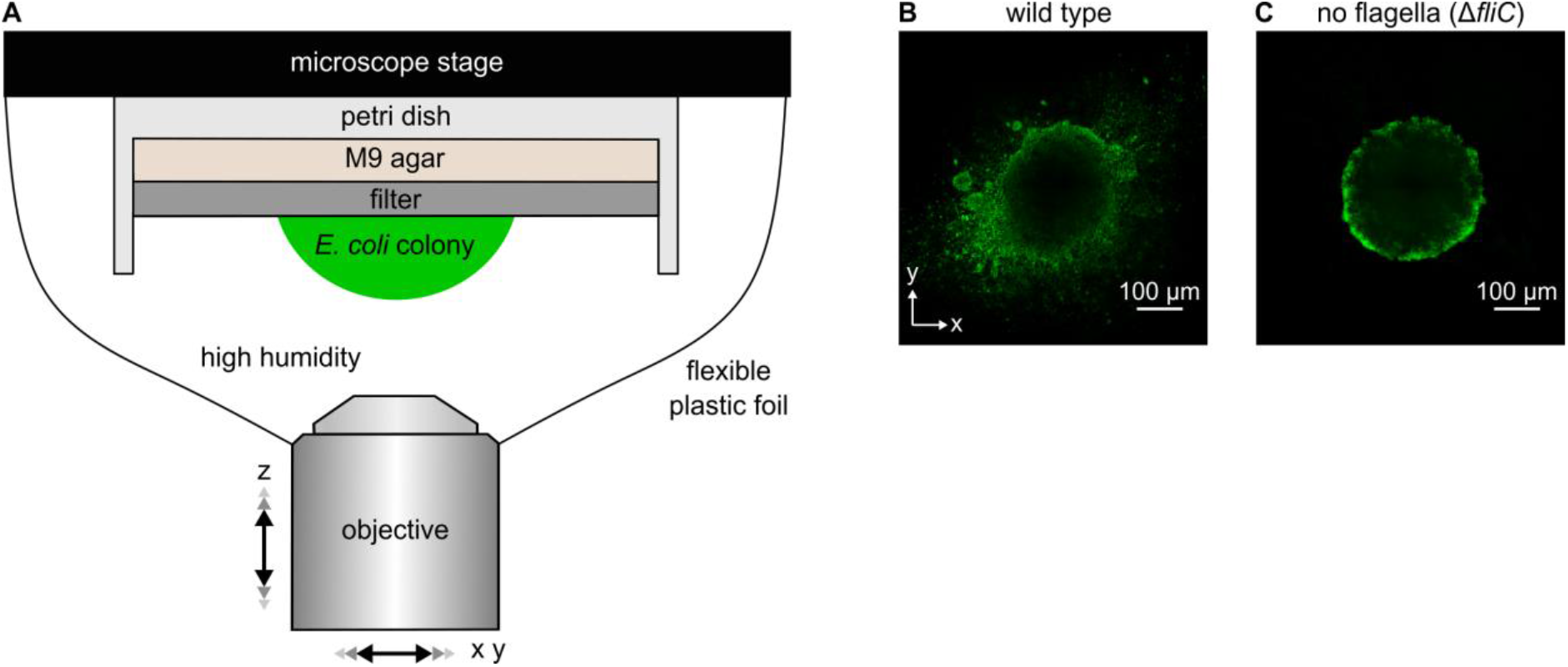
In contrast to wild-type colonies, flagella-deficient (Δ*fliC*) colonies are highly symmetric in shape. (**A**) Experimental arrangement for investigating colony growth using an inverted confocal microscope. Dimensions are not drawn to scale. (**B**, **C**) Confocal microscopy images of *xy*-planes of colonies on filter membranes on M9 agar, made by the wild type strain (panel **B**) or a strain lacking the flagellin gene *fliC* (panel **C**) grown for 24 h. The Δ*fliC* strain produces no flagella and is incapable of swimming motility.

**Figure 1—figure supplement 2.**
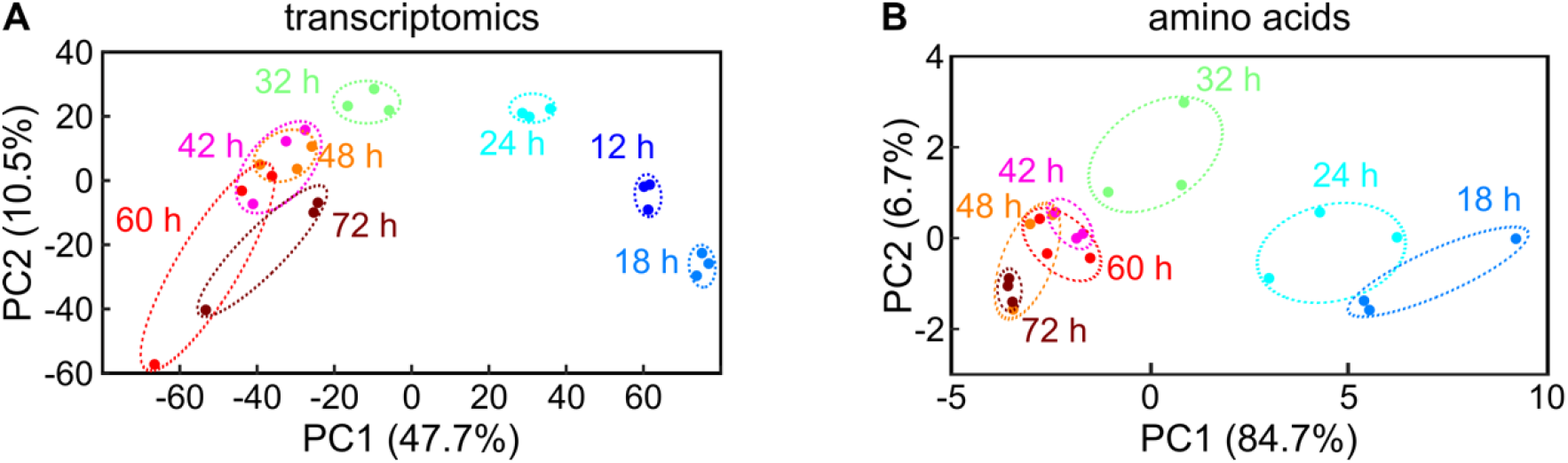
Principal component analysis of transcriptomic and metabolomic data. (**A**) Principal component analysis of the transcriptomic data presented in Figure 1C. The axes represent the 1^st^ principal component (PC1) and the 2^nd^ principal component (PC2). (**B**) Principal component analysis of amino acid data presented in Figure 1E. Each circle represents a different independent transcriptome or metabolome replicate. Replicates for the same colony growth timepoint have the same color and are enclosed by an ellipse with a dashed line.

**Figure 1—figure supplement 3.**
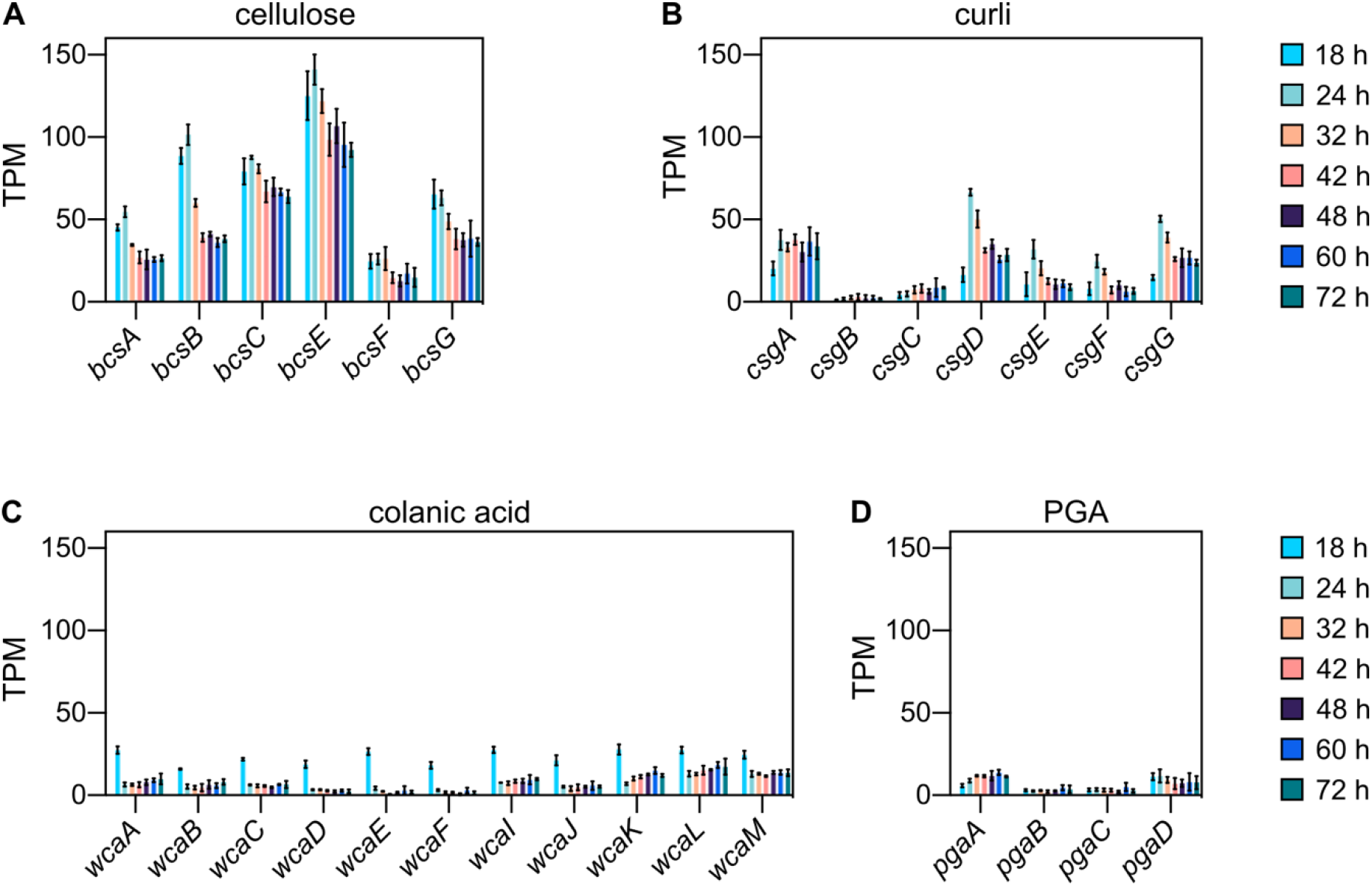
*E. coli* biofilm colonies express extracellular matrix genes. *E. coli* extracellular matrix is composed of cellulose, curli, colanic acid, *β*-1,6-N-acetyl-D-glucosamine (PGA) and flagella (Hobley et al., 2015). Here, transcripts per million (TPM) of genes involved in the production of cellulose (**A**), curli (**B**), colonic acid (**C**) and PGA (**D**) are shown for colonies of the parental strain (KDE722) grown for 18 h, 24 h, 32 h, 42 h, 48 h, 60 h, or 72 h, measured using RNA-seq. Flagellar genes are not shown here, because the flagellin gene *fliC* is deleted in the parental strain. Data are mean ± standard deviation (s.d.), *n* = 3.

**Figure 1—figure supplement 4.**
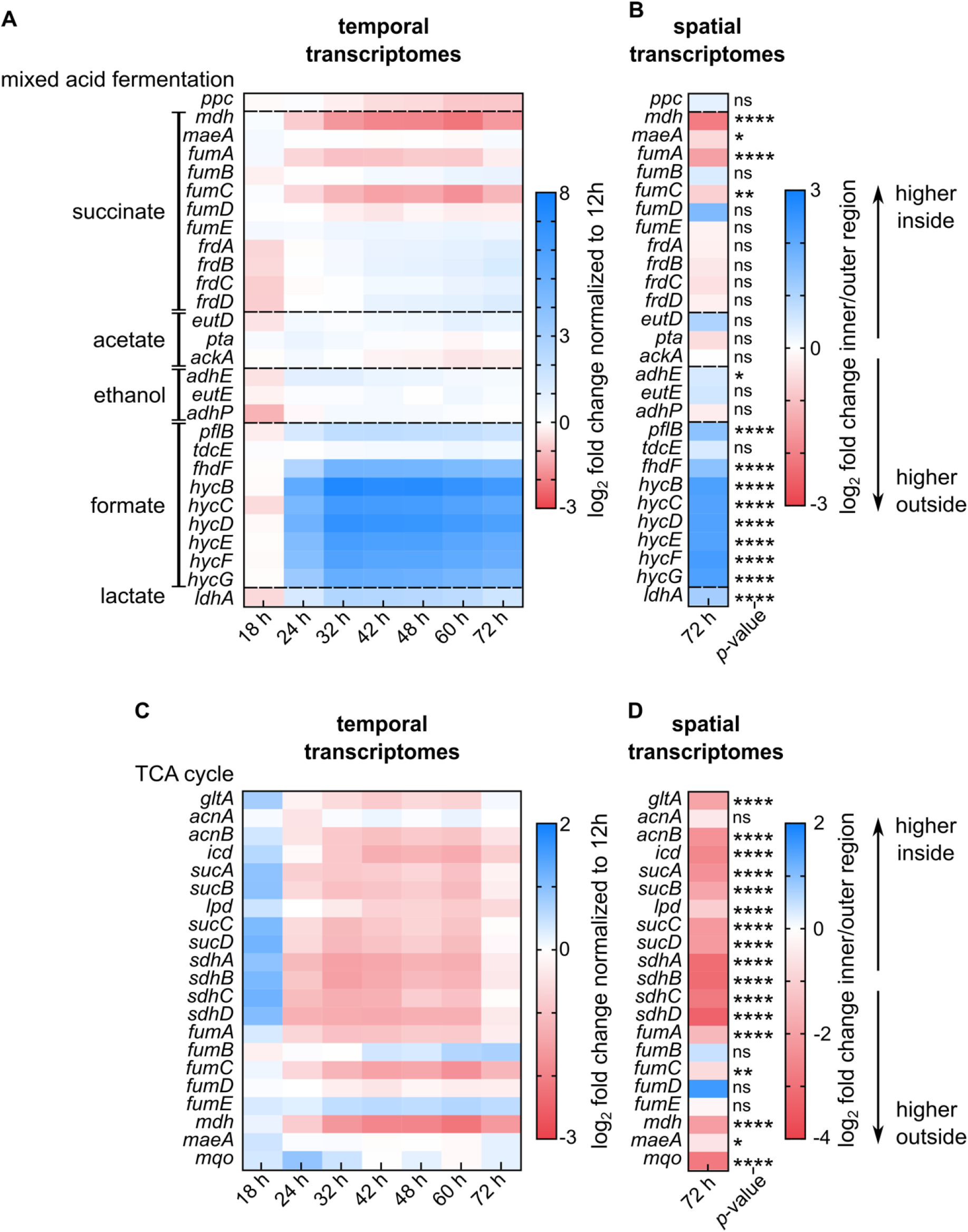
Spatial and temporal expression of genes involved in mixed acid fermentation and TCA cycle in colonies on M9 agar. (**A**, **B**) Heatmaps show genes involved in mixed acid fermentation. (**A**) Fold-changes of genes from whole-colony measurements at different growth stages, normalized to transcriptomes of 12 h colonies. (**B**) Fold-changes of genes from spatially-resolved transcriptomes, measured for 72 h colonies. The inner region corresponds to non-fluorescent cells, the outer region corresponds to those cells with mRuby2 fluorescence. (**C**, **D**) Heatmaps show genes involved in the TCA cycle. (**C**) Fold-changes of genes from whole-colony measurements at different growth stages, normalized to transcriptomes of 12 h colonies. (**D**) Fold-changes of genes from spatially-resolved transcriptomes, measured for 72 h colonies. Non-significant differences are labelled “ns”. *p*-values correspond to FDR-adjusted *p*-values: * *p* < 0.05, ** *p* < 0.01, **** *p* < 0.0001. For panels *A* and *C*, *n* = 3; for *B* and *D*, *n* = 4. Some genes are involved in both pathways and are therefore displayed in all heatmaps.

**Figure 1—figure supplement 5.**
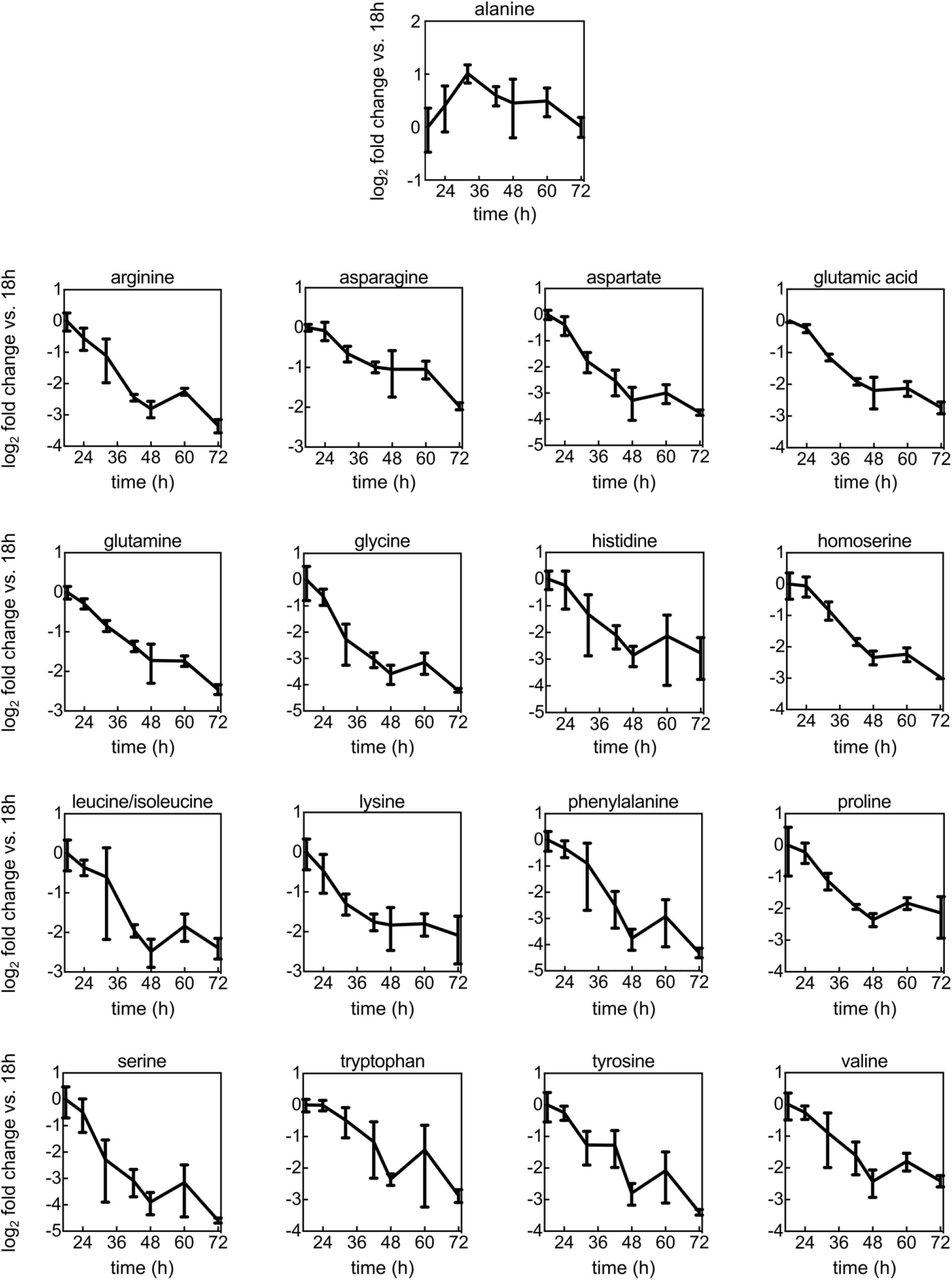
Amino acid levels during colony growth. Fold-changes of amino acid levels as a function of time during colony development on M9 agar, normalized to amino acid levels from 18 h colonies. Measurements were performed using mass spectrometry. Data are mean values ± s.d., *n* = 3.

**Figure 2—figure supplement 1.**
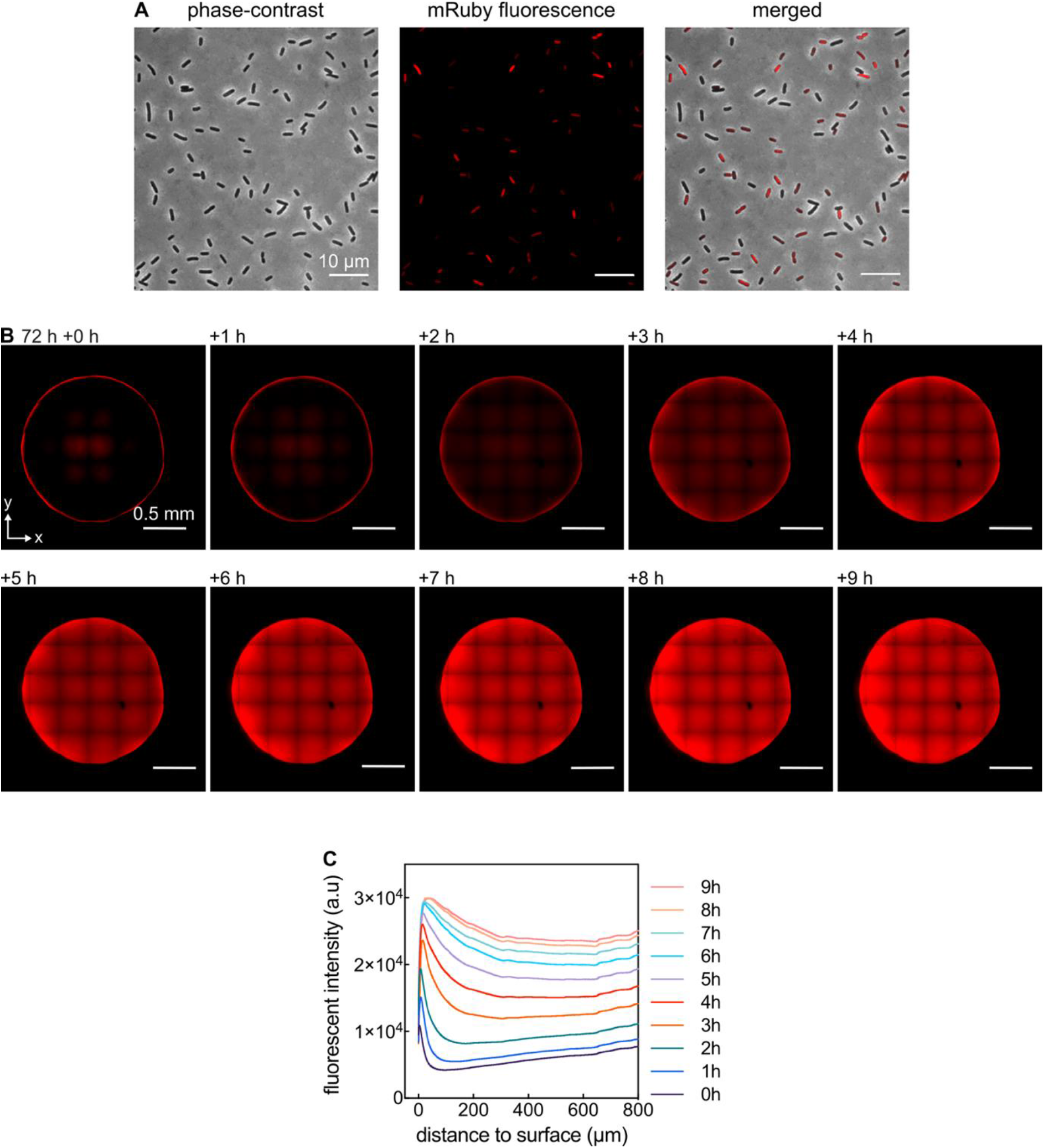
Fluorescent gradients of mRuby2 in colonies are not due to imaging artefacts. (**A**) Images of cells from a 72 h colony that was resuspended in PBS. Some cells are fluorescent, others display less or no fluorescence. (**B**) Confocal images of a *xy*-plane of a 72 h colony acquired at *z* = 60 μm above the filter membrane on M9 agar. At 72 h + 0 h the filter membrane carrying the colony was moved from M9 agar to an agar plate with the same medium, but lacking glucose. As a consequence, cells stop growth and stop consuming molecular oxygen, allowing oxygen to penetrate into the colony. Penetration of oxygen allows the mRuby2 protein to fold into the fluorescent conformation. The square pattern in the images results from the fact that colony images were stitched together from 7 × 7 image tiles. (**C**) Quantification of panel **B**: Fluorescent intensity for different times after transferring the colony to M9 agar without glucose (different colored lines), as a function of distance to the outer surface of the colony.

**Figure 2—figure supplement 2.**
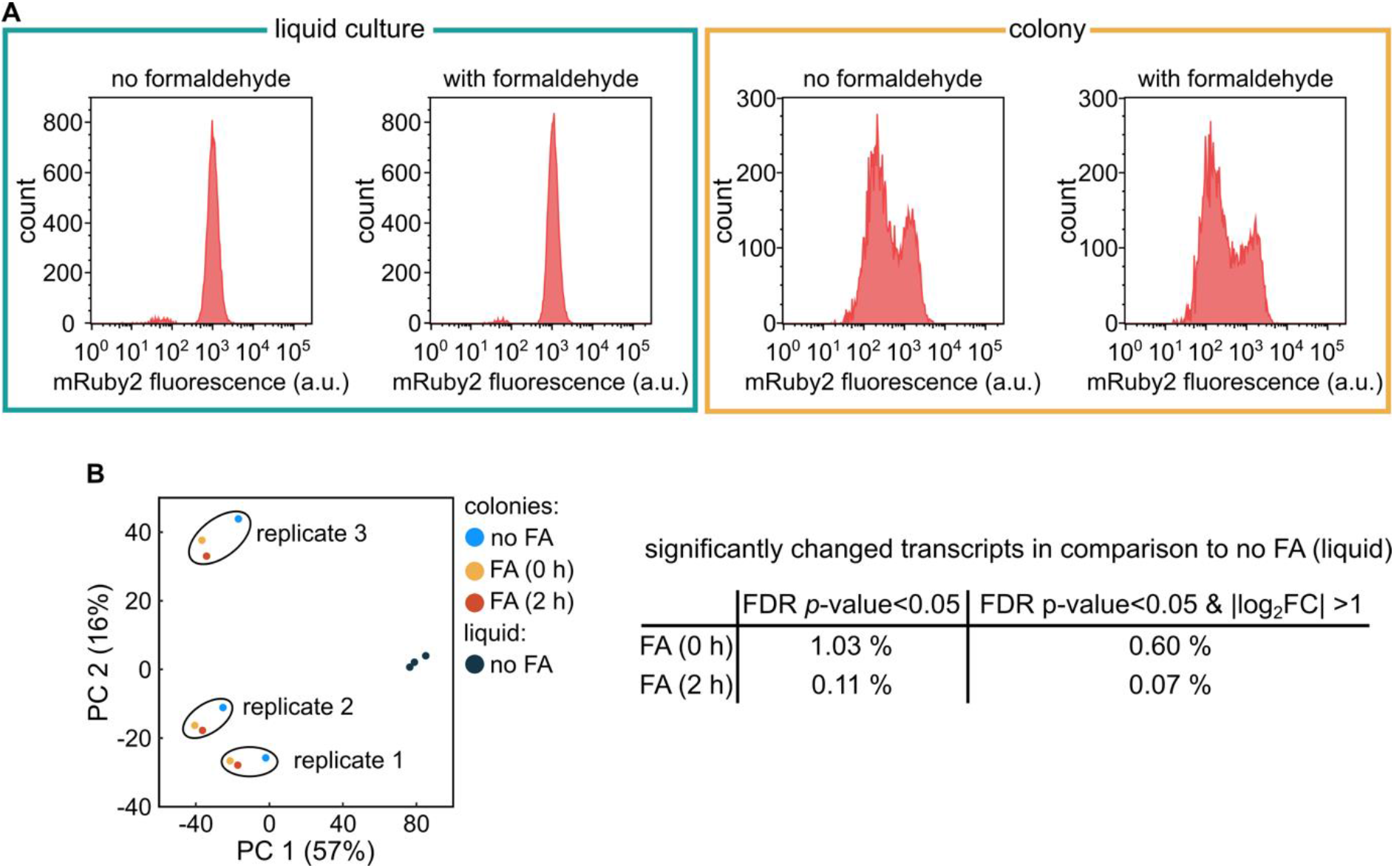
Formaldehyde fixation does not affect mRuby2 fluorescence or the colony transcriptome. (**A**) Flow cytometry results for fluorescence of *E. coli* constitutively expressing mRuby2 (strain KDE722) with or without 2 hours of fixation using 4% formaldehyde. For the two panels on the left (enclosed by green box), the cells were grown in liquid shaking M9 medium. For the two panels on the right (enclosed by orange box), the cells were grown as a colony on M9 agar. (**B**) Principal component analysis of the transcriptomes of colonies grown for 72 h, which were either fixed with 4% formaldehyde (FA) for 0 h or 2 h, or without fixation. As a control, the transcriptome of cells from an exponentially growing M9 liquid shaking culture without fixation was also analyzed. Each data point corresponds to an independent replicate (*n* = 3). The table on the right indicates the number of differentially expressed genes in colonies, in comparison to the colonies without fixation, using as threshold for differentially expressed genes either a FDR-adjusted *p*-value < 0.05, or a FDR-adjusted *p*-value < 0.05 and an absolute log_2_ fold-change (log_2_ FC) > 1.

**Figure 2—figure supplement 3.**
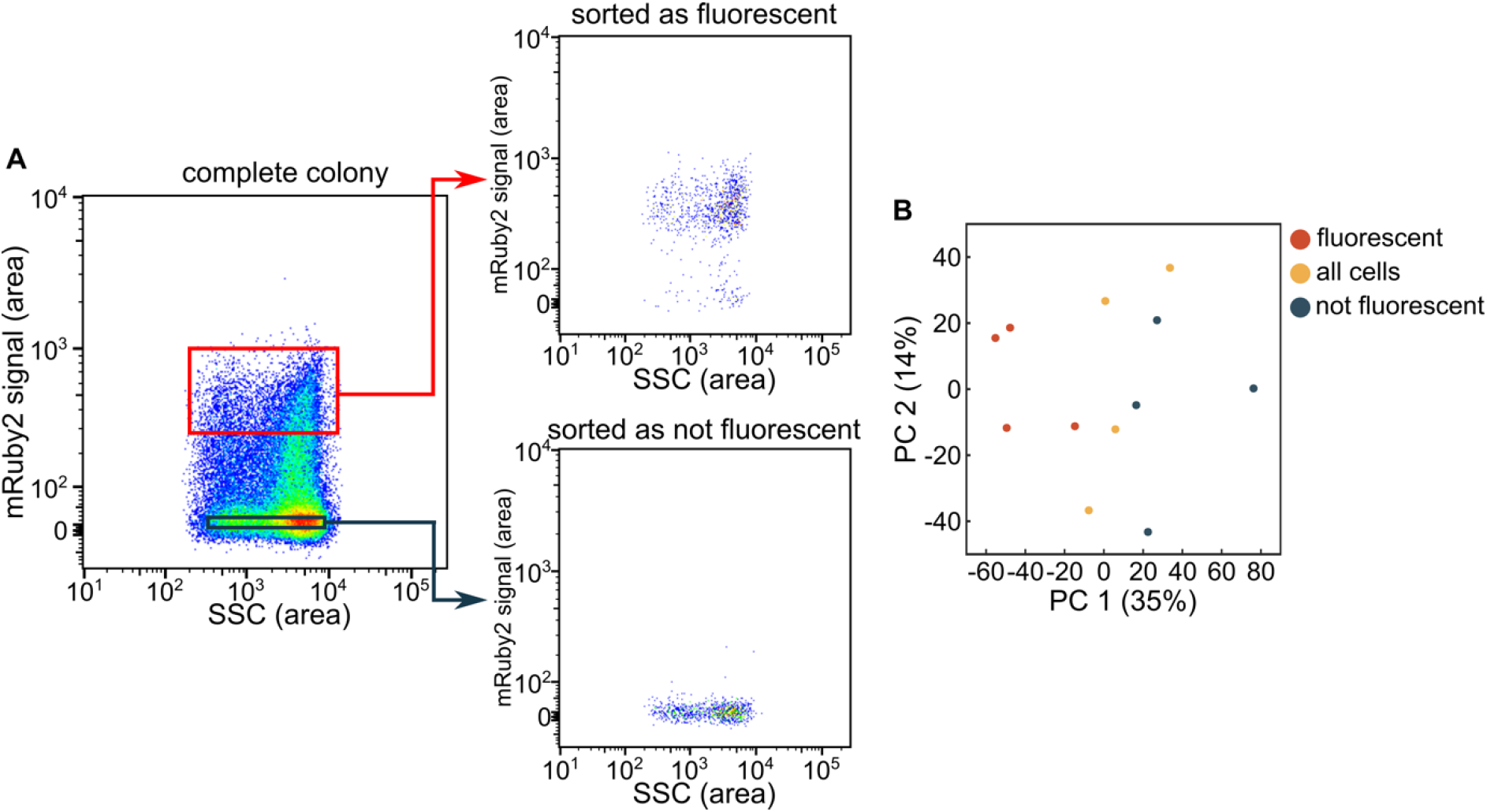
Application of flow cytometry with fluorescence-activated cell sorting to separate cells from the inner and outer regions of colonies. (**A**) Representative sorting results of a colony grown for 72 h, using the mRuby2 fluorescence signal and the side scatter (SSC) signal. Cells were sorted into two separate bins (red, black). (**B**) Principal component analysis of RNA-seq results of the two separated bins, or the entire population, from colonies that were grown for 72 h, showing the 1^st^ (PC1) and 2^nd^ (PC2) principal component axes. Each data point corresponds to an independent replicate (*n* = 4).

**Figure 2—figure supplement 4.**
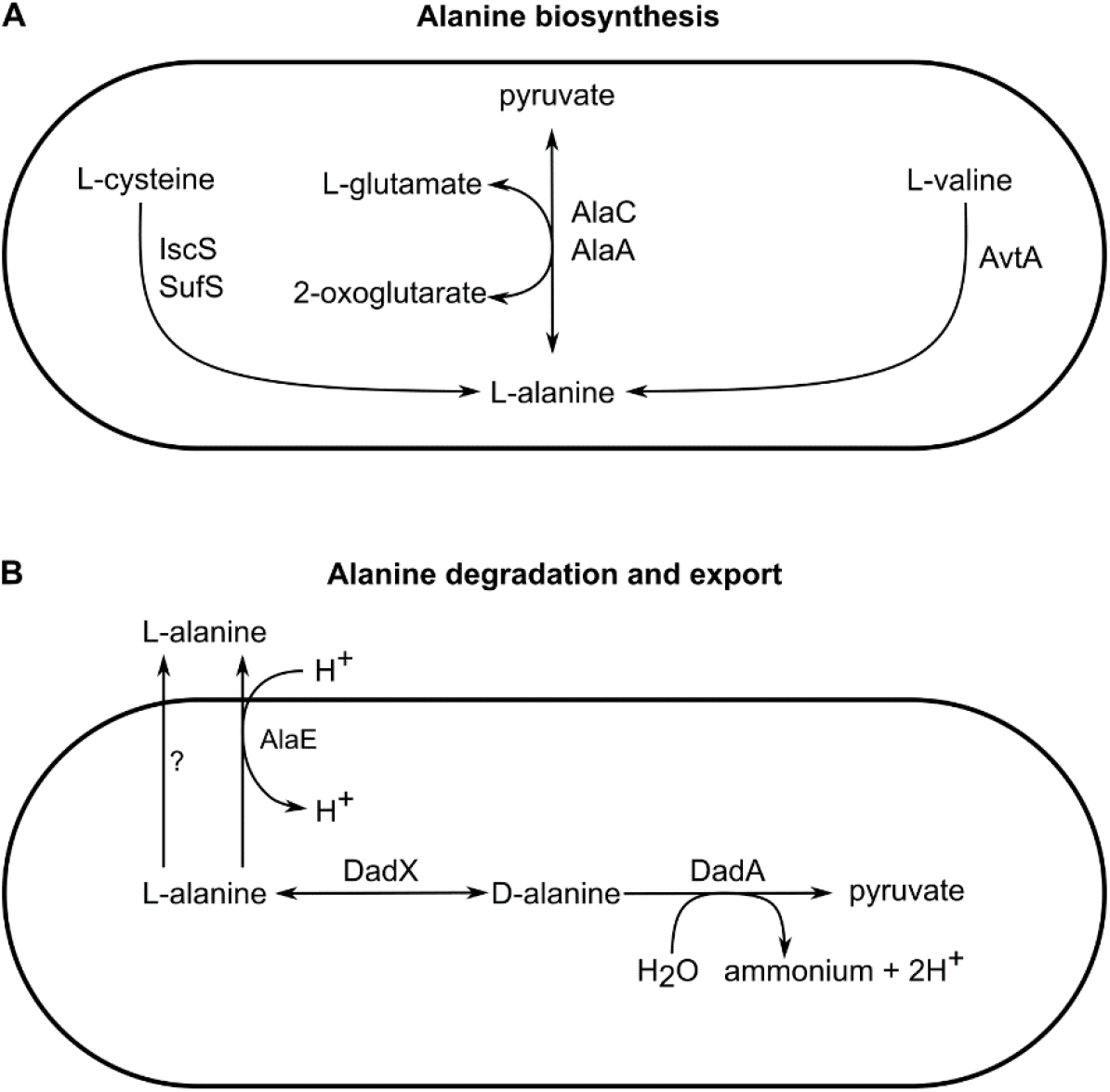
Key pathways in alanine metabolism. (**A**) Alanine biosynthesis pathways. (**B**) Alanine degradation and export pathways. Cells are drawn with a schematic rod shape.

**Figure 4—figure supplement 1.**
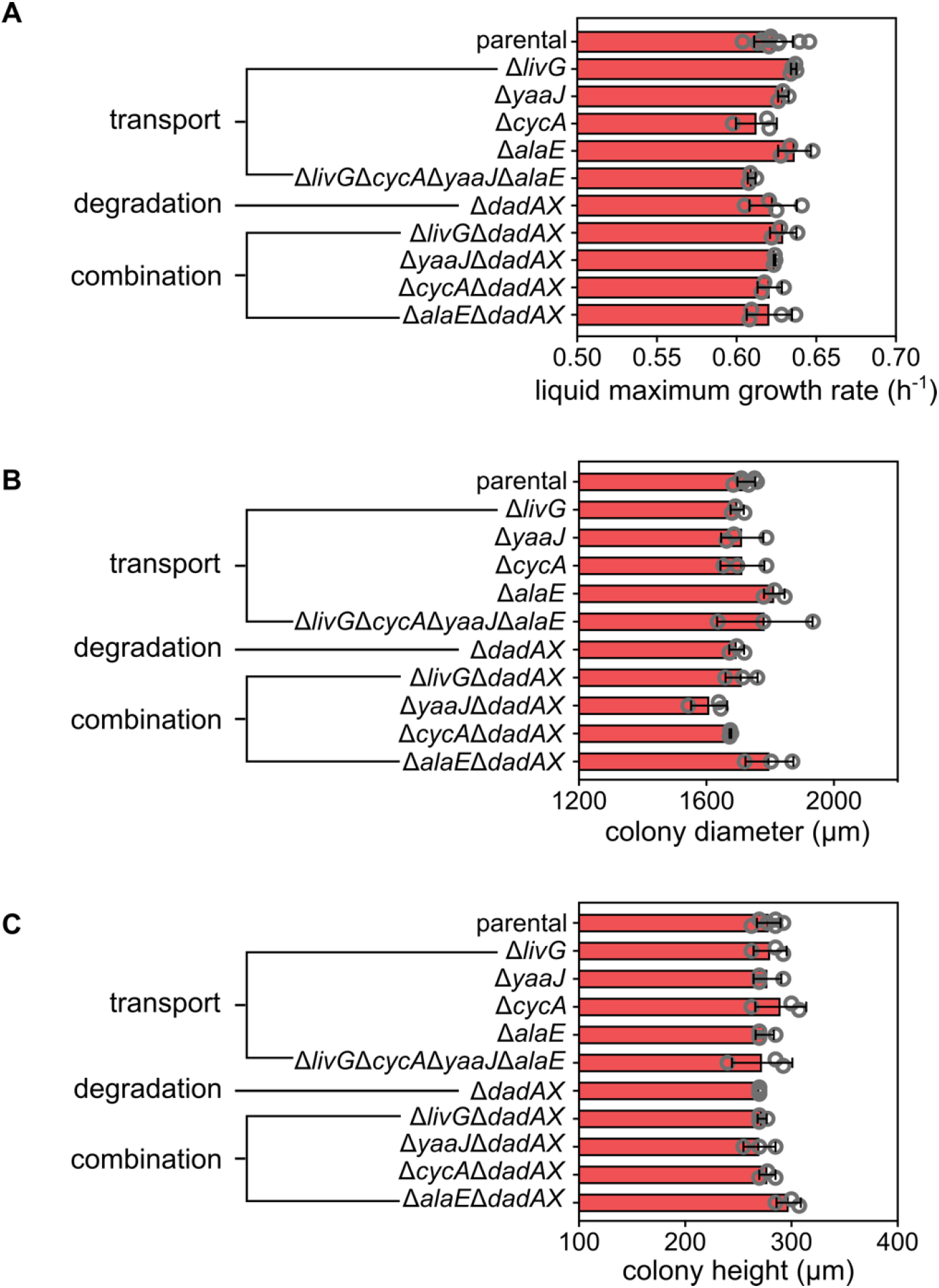
Growth rates and colony sizes for alanine metabolism mutants. (**A**) Maximum growth rate in liquid shaking cultures in M9 medium. Colony diameter (**B**) and height (**C**) after 72 h of growth on M9 agar for different mutants in alanine transport and degradation, compared to the parental *E. coli* strain. Data are mean values ± s.d., each data point is shown as a grey circle.

**Figure 4—figure supplement 2.**
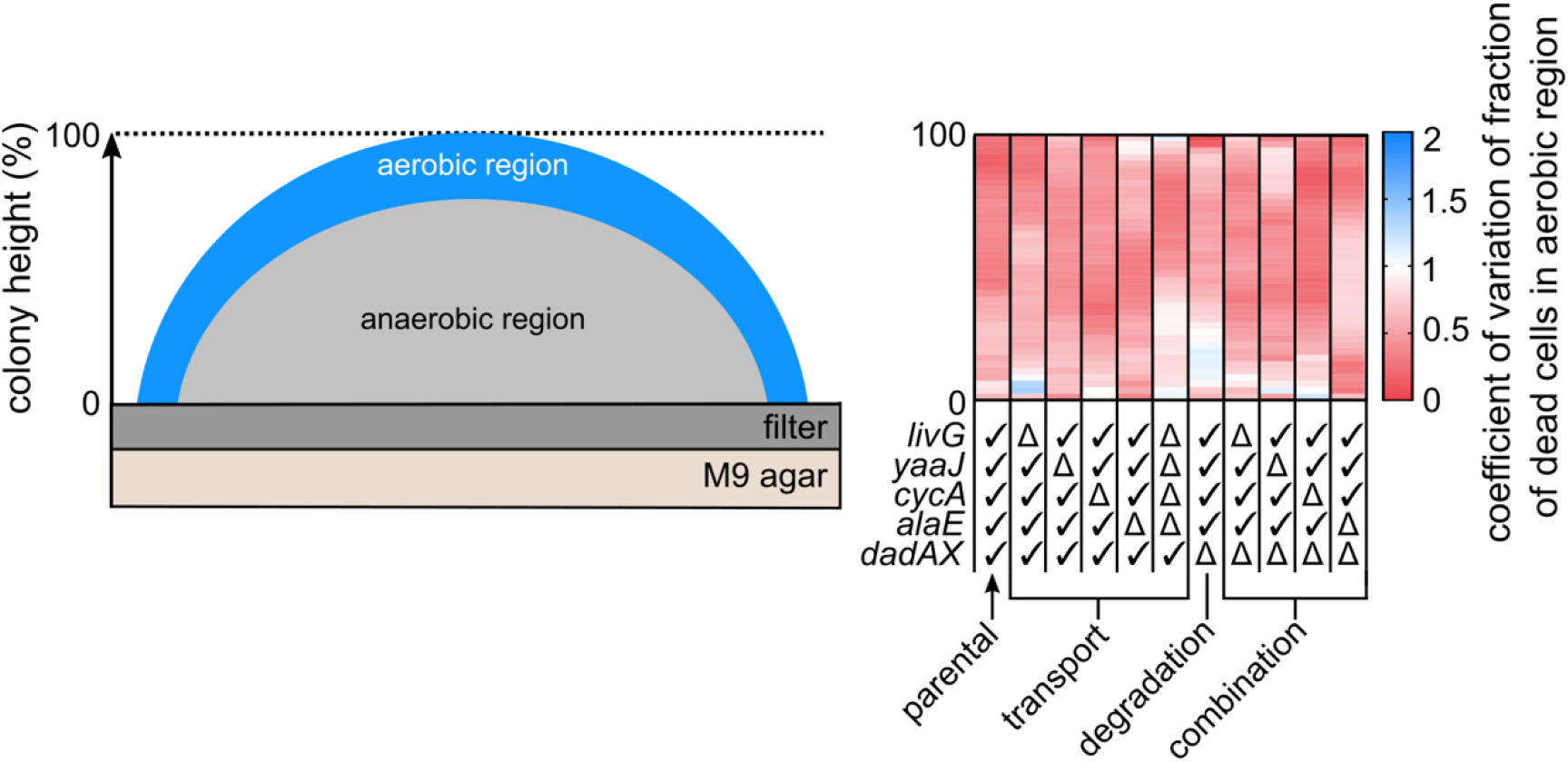
Coefficient of variation of the fraction of dead cells in the aerobic region of the colony. Heatmap shows the coefficient of variation for the data shown in Figure 4A.

**Figure 4—figure supplement 3.**
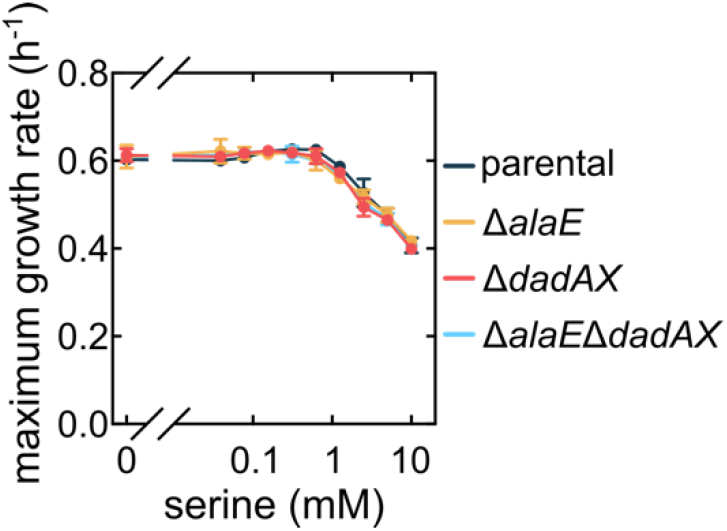
Effects of exogenously added alanine on maximum growth rate for alanine metabolism mutants are specific to alanine, and are not triggered serine. The maximum liquid culture growth rate as function of the concentration of exogenically added serine in M9 medium is the same for all strains. Data are mean values ± s.d., *n* = 3.

**Figure 5—figure supplement 1.**
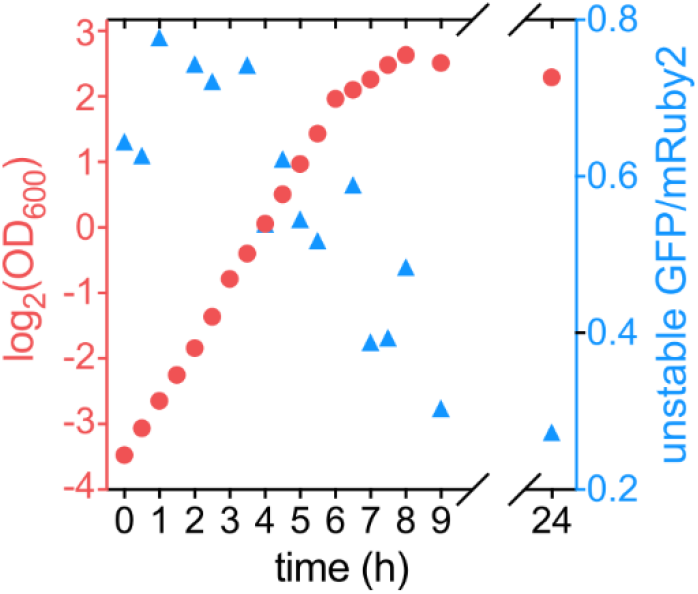
Ratio between unstable GFP and stable mRuby2 correlates with bacterial growth rate. Optical density at 600 nm (OD_600_; shown in red, left y-axis) and the ratio between unstable GFP (with the ASV-tag) and mRuby2 (shown in blue, right y-axis) as a function of time for a liquid culture of strain KDE2937.

**Table S1:** Bacterial strains used in this study. Abbreviations: Kan = kanamycin. Superscript “R” = resistance. “-” = fusion. “::” = insertion. The scar corresponds to 5’-GAAGTTCCTATACTTTCTAGAGAATAGGAACTTC-3’ sequence.

| Strain | Genotype/ Relevant features | Reference |
| --- | --- | --- |
| KDE261 | *E. coli* strain carrying plasmid pCP20. | Drescher lab stock |
| KDE262 | *E. coli* strain carrying plasmid pKD46. | Drescher lab stock |
| KDE264 | *E. coli* strain carrying plasmid pKD3. | Drescher lab stock |
| KDE265 | *E. coli* strain carrying plasmid pKD4. | Drescher lab stock |
| KDE1361 | *E. coli* strain carrying plasmid pNUT1361. | Drescher lab stock |
| KDE2338 | *E. coli* strain carrying plasmid pNUT2338. | Drescher lab stock |
| KDE2658 | *E. coli* strain carrying plasmid pUC18R6KT-mini-Tn7-Km (Addgene #64969). | Drescher lab stock |
| KDE2659 | *E. coli* strain carrying plasmid pTNS2 (Addgene #64968). | Drescher lab stock |
| KDE2674 | *E. coli* strain carrying plasmid pNUT2674. | This study |
| KDE2787 | *E. coli* strain carrying plasmid pNUT2787. | This study |
| KDE2838 | *E. coli* strain carrying plasmid pNUT2838. | This study |
| KDE474 | *E. coli* AR3110 WT. | (Serra, Richter, & Hengge, 2013) |
| KDE679 | AR3110, *P<sub>tac</sub>-mRuby2-mRuby2* and *Kan<sup>R</sup>* inserted at *attB* site (*P<sub>tac</sub>* without operator). | (Vidakovic et al., 2018) |
| KDE722 | KDE679 with $\Delta fliC::scar$. | (Vidakovic et al., 2018) |
| KDE1899 | AR3110 with $\Delta fliC::scar$. | This study |
| KDE2007 | KDE679 with $\Delta fliC::scar$, $\Delta alaE::scar$. | This study |
| KDE2009 | KDE679 with $\Delta fliC::scar$, $\Delta dadAX::scar$. | This study |
| KDE2086 | KDE679 with $\Delta fliC::scar$, $\Delta alaE::scar$, $\Delta dadAX::scar$. | This study |
| KDE2183 | KDE679 with $\Delta fliC::scar$, $\Delta cycA::scar$. | This study |
| KDE2185 | KDE679 with $\Delta fliC::scar$, $\Delta livG::scar$. | This study |
| KDE2438 | KDE679 with $\Delta fliC::scar$, $\Delta yaaJ::scar$. | This study |
| KDE2533 | KDE679 with $\Delta fliC::scar$, $\Delta cycA::scar$, $\Delta livG::scar$, $\Delta alaE::scar$, $\Delta yaaJ::scar$. | This study |
| KDE2564 | KDE679 with $\Delta fliC::scar$, $\Delta cycA::scar$, $\Delta dadAX::scar$. | This study |
| KDE2607 | KDE679 with $\Delta fliC::scar$, $\Delta yaaJ::scar$, $\Delta dadAX::scar$. | This study |
| KDE2937 | KDE679 with $\Delta fliC::scar$, *P<sub>tac</sub>-sfgfp*(ASV) at the Tn7 insertion site, coding for an unstable superfolder GFP with an AANDENYAASV-tag. | This study |
| KDE2938 | KDE679 with $\Delta fliC::scar$, $\Delta alaE::scar$, *P<sub>tac</sub>-sfgfp*(ASV) at the Tn7 insertion site. | This study |
| KDE2939 | KDE679 with $\Delta fliC::scar$, $\Delta dadAX::scar$, *P<sub>tac</sub>-sfgfp*(ASV) at the Tn7 insertion site. | This study |
| KDE2940 | KDE679 with $\Delta fliC::scar$, $\Delta alaE$ $\Delta dadAX$, *P<sub>tac</sub>-sfgfp*(ASV) at the Tn7 insertion site. | This study |

**Table S2:**
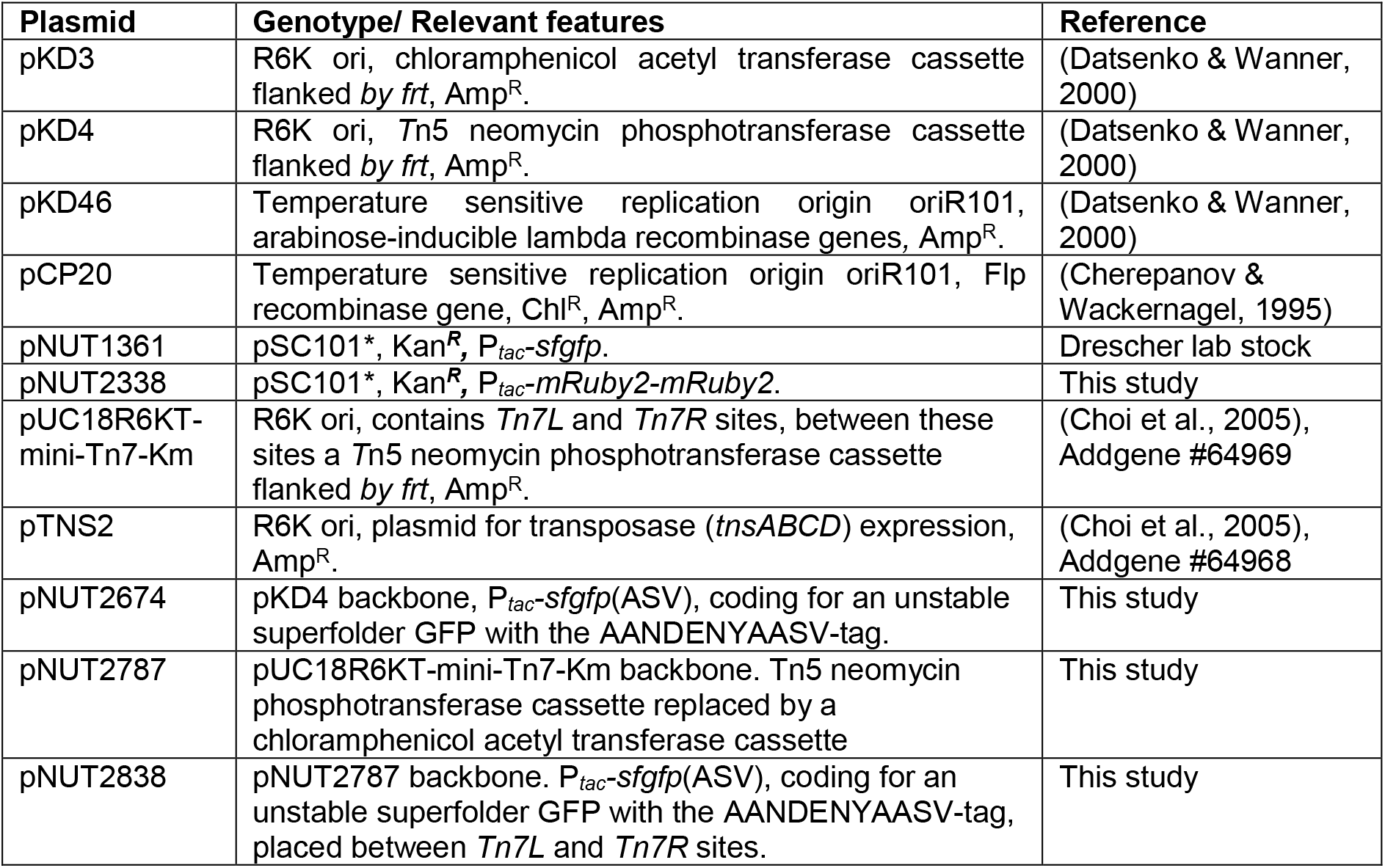
Plasmids used in this study. Abbreviations: Kan = kanamycin, Amp = ampicillin, Chl = chloramphenicol. Superscript “R” = resistance. “-“ = fusion.

| Plasmid | Genotype/ Relevant features | Reference |
| --- | --- | --- |
| pKD3 | R6K ori, chloramphenicol acetyl transferase cassette flanked by *frt*, Amp<sup>R</sup>. | (Datsenko & Wanner, 2000) |
| pKD4 | R6K ori, *Tn5* neomycin phosphotransferase cassette flanked by *frt*, Amp<sup>R</sup>. | (Datsenko & Wanner, 2000) |
| pKD46 | Temperature sensitive replication origin oriR101, arabinose-inducible lambda recombinase genes, Amp<sup>R</sup>. | (Datsenko & Wanner, 2000) |
| pCP20 | Temperature sensitive replication origin oriR101, Flp recombinase gene, Chl<sup>R</sup>, Amp<sup>R</sup>. | (Cherepanov & Wackernagel, 1995) |
| pNUT1361 | pSC101*, Kan<sup>R</sup>, P<sub>tac</sub>-*sfgfp*. | Drescher lab stock |
| pNUT2338 | pSC101*, Kan<sup>R</sup>, P<sub>tac</sub>-*mRuby2-mRuby2*. | This study |
| pUC18R6KT-mini-Tn7-Km | R6K ori, contains *Tn7L* and *Tn7R* sites, between these sites a *Tn5* neomycin phosphotransferase cassette flanked by *frt*, Amp<sup>R</sup>. | (Choi et al., 2005), Addgene #64969 |
| pTNS2 | R6K ori, plasmid for transposase (*tnsABCD*) expression, Amp<sup>R</sup>. | (Choi et al., 2005), Addgene #64968 |
| pNUT2674 | pKD4 backbone, P<sub>tac</sub>-*sfgfp*(ASV), coding for an unstable superfolder GFP with the AANDENYAASV-tag. | This study |
| pNUT2787 | pUC18R6KT-mini-Tn7-Km backbone. *Tn5* neomycin phosphotransferase cassette replaced by a chloramphenicol acetyl transferase cassette | This study |
| pNUT2838 | pNUT2787 backbone. P<sub>tac</sub>-*sfgfp*(ASV), coding for an unstable superfolder GFP with the AANDENYAASV-tag, placed between *Tn7L* and *Tn7R* sites. | This study |

**Table S3:**
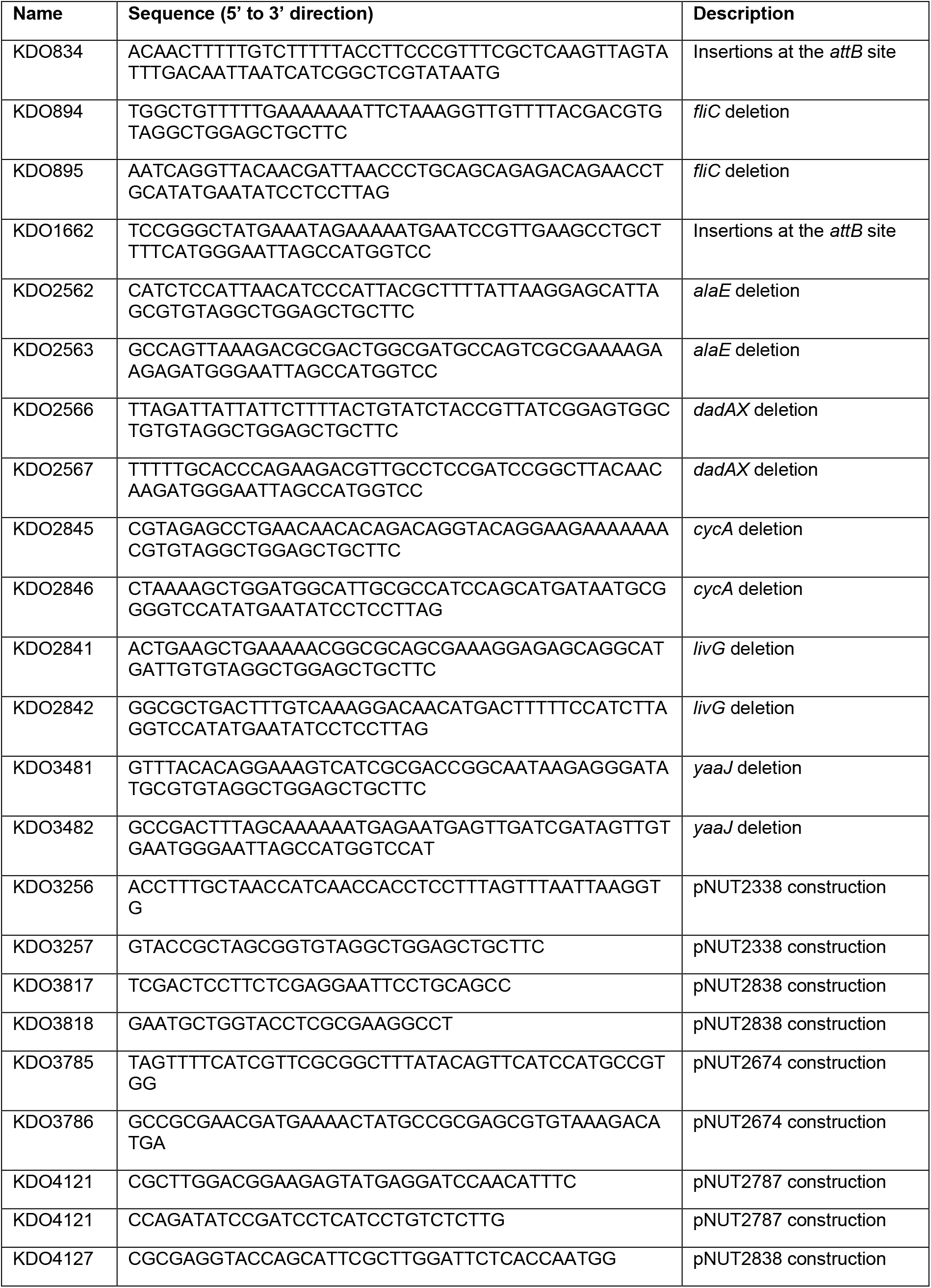
DNA oligonucleotides used in this study.

